# Herpes zoster mRNA vaccine induces superior vaccine immunity over licensed vaccine in mice and rhesus macaques

**DOI:** 10.1101/2023.08.16.553640

**Authors:** Lulu Huang, Tongyi Zhao, Weijun Zhao, Andong Shao, Huajun Zhao, Wenxuan Ma, Yingfei Gong, Xianhuan Zeng, Changzhen Weng, Lingling Bu, Zhenhua Di, Shiyu Sun, Qinsheng Dai, Minhui Sun, Limei Wang, Zhenguang Liu, Leilei Shi, Jiesen Hu, Shentong Fang, Cheng Zhang, Jian Zhang, Guan Wang, Karin Loré, Yong Yang, Ang Lin

## Abstract

Herpes zoster remains an important global health issue and mainly occurs in aged and immunocompromised individuals with an early exposure history to Varicella Zoster Virus (VZV). Although the licensed vaccine Shingrix has a remarkably high efficacy, undesired reactogenicity and increasing global demand causing vaccine shortage urged the development of improved or novel VZV vaccines. In this study, we developed a novel VZV mRNA vaccine candidate (named as ZOSAL) containing sequence-optimized mRNAs encoding full-length glycoprotein E encapsulated in an ionizable lipid nanoparticle. In mice and rhesus macaques, ZOSAL demonstrated superior immunogenicity and safety in multiple aspects over Shingrix, especially in the induction of strong T cell immunity. Transcriptomic analysis revealed that both ZOSAL and Shingrix could robustly activate innate immune compartments, especially Type-I IFN signaling and antigen processing/presentation. Multivariate correlation analysis further identified several early factors of innate compartments that can predict the magnitude of T cell responses, which further increased our understanding of the mode of action of two different VZV vaccine modalities. Collectively, our data demonstrated the superiority of VZV mRNA vaccine over licensed subunit vaccine. The mRNA platform therefore holds prospects for further investigations in next-generation VZV vaccine development.

## Introduction

Varicella Zoster Virus (VZV) is a human alphaherpesvirus that causes varicella during primary infection and establishes latency in the sensory ganglia posing the risks for reactivation to cause herpes zoster (HZ) [1]. HZ mainly occurs in aged and immunocompromised individuals with an early exposure history and is typically symptomized by a painful dermatomal rash or more seriously postherpetic neuralgia (PHN). The genome of VZV encodes eight glycoproteins that are critical for viral entry and replication [1,2]. Amongst them, glycoprotein E (gE) is a 623-aa transmembrane protein containing a 544-aa hydrophilic ectodomain, a 17-aa hydrophobic transmembrane region and a 62-aa cytoplasmic region. gE is abundantly present on the surface of viral particle and also on the plasma membrane of VZV-infected cells, which has been reported to mediate viral spread and skin tropism, as well as the formation of infectious virions [1]. Moreover, gE is highly immunogenic in eliciting both antibody and cell-mediated immune (CMI) responses and is believed to be a key immunogen for VZV vaccine development [2–4]. A live-attenuated vaccine (Zostavax, Merck) and a protein-based subunit vaccine containing carboxyl-terminal truncated form of gE adjuvanted with AS01B (Shingrix) have been approved in clinical use, of which Shingrix showed a remarkably higher protective efficacy (97.2%) than Zostavax (51.3%) in adults above 50 years of age [5]. However, high reactogenicity with both local and systemic reaction and the increasing global demand causing vaccine shortage remain major concerns with Shingrix, which warrant further development of improved or novel VZV vaccines [6–9]. Recently, a domestic live-attenuated VZV vaccine developed by Changchun Bcht Biotech was approved for adult use in China, but details on the protective efficacy of this vaccine was yet undisclosed.

VZV-specific T cell responses are critical in the prevention and control of initial VZV infection, as well as reactivation of latent infection [10–12]. Clinical evidence has demonstrated that higher levels of VZV-specific T cells, not specific antibodies (Abs), are associated with reduced HZ incidence and disease severity [13]. More in-depth studies using simian varicella virus (SVV)-infected rhesus macaques that recapitulate VZV infection in humans showed that depletion of CD4^+^ T cells led to prolonged viremia and severe varicella during SVV infection. While in contrast, depletion of CD20^+^ B cells or CD8^+^ T cells did not alter the severity of varicella [14]. Further depletion of CD4^+^ cells in rhesus macaques with pre-established SVV latency led to viral reactivation causing zoster rash [15]. In addition, lower frequencies of VZV-specific IFN-γ and TNF-producing CD4^+^ T cells, not CD8^+^ T cells, were found to be associated with higher incidence of HZ in patients with systemic lupus erythematosus [16]. All these suggested that there is a more central role of VZV-specific CD4^+^ T cells rather than CD8^+^ T cells or Abs in controlling VZV infection. To this end, strong virus-specific T cell responses, particularly the Th1-type CD4^+^ T cells, are indispensable to be induced by novel VZV vaccine candidates. The mRNA platform has shown multiple advantages in vaccine development including its ability to elicit strong T cell responses, which may largely be attributed to the innate immune stimulatory effects of both mRNA cargo and lipid components in the delivery system [17–20]. A novel VZV mRNA vaccine candidate from Moderna Therapeutics has recently been tested when given as a booster dose in rhesus macaques that had been prime immunized with a live-attenuated VZV vaccine and was shown to induce comparable levels of Abs and Th1-biased T cells to Shingrix [21]. This suggested a high potentiality of mRNA technology in the development of next-generation VZV vaccines, which awaits further investigations.

In the current study, we developed a novel VZV mRNA vaccine (named as ZOSAL) containing sequence-optimized mRNAs encoding full-length glycoprotein E (gE) encapsulated in an ionizable lipid nanoparticle (LNP) formulation using our well-established platforms [22–24]. The immunogenicity of ZOSAL was thoroughly evaluated and compared side-by-side with the benchmark vaccine (Shingrix) in adult mice, aged mice, and importantly in rhesus macaques. To better understand the generation of vaccine-induced VZV-specific immunity and to elucidate the differences between two vaccine modalities, we performed in-depth characterization of the vaccine responses generated in rhesus macaques, with particular emphasis on the innate immune gene signatures and magnitude and functionality of B cell and T cell responses. Apart from these, some of the key safety aspects of ZOSAL and Shingrix were assessed by longitudinally monitoring the hematological and serum biochemical parameters, as well as the early changes in safety-related genes. Our data showed that, in multiple animal models, ZOSAL elicited superior VZV-specific immunity and meanwhile demonstrated higher safety profiles over Shingrix. We also identified correlates of vaccine immunity that may impact on the development of vaccine responses, which further increases our understanding of the mode of action of VZV vaccines.

## Materials and Methods

A detailed description and additional materials and methods are available in supplemental materials.

### Ethics, animals, immunization

All animal experiments were performed in accordance with the Guidelines for the Care and Use of Laboratory Animals and the Ethical Committee of China Pharmaceutical University and using protocols approved by the Institutional Animal Care and Use Committee of China Pharmaceutical University (Approval number: AP-B2209P011). C57BL/6 mice (female, 6-week or 10-month old) were purchased from Hangzhou Ziyuan Laboratory Animal Technology Co., Ltd. and randomly allocated to different groups. Mice were immunized intramuscularly (i.m.) twice at an interval of 2 weeks. Sera samples were collected longitudinally for Ab response analysis. Mice were necropsied at the indicated time points after vaccination. Spleens and draining lymph nodes (dLN) were collected and processed to obtain single cell suspension for analysis. For the nonhuman primate (NHP) experiments, Chinese rhesus macaques (3-5 years old) were housed at the Center for New Drug Safety Evaluation and Research at China Pharmaceutical University. Animals were i.m. immunized with ZOSAL (100μg per dose) or Shingrix (human dose) at day 0 and 28. Peripheral venous blood was collected in EDTA-2K tubes and processed within 2 hours to obtain fresh peripheral blood mononuclear cells (PBMCs) according to the previously reported protocol [25]. Serum samples and PBMCs were collected longitudinally at different time points before or after vaccination. Body weight, temperature, blood cell count and biochemical parameters were monitored longitudinally for analysis of safety profiles.

### mRNA vaccine preparation

VZV mRNA vaccine (ZOSAL) was prepared as previously described [22,23]. In brief, mRNA encoding for VZV gE immunogen was synthesized in vitro by T7 polymerase-mediated transcription from a linearized DNA template. Methyl-pseudouridine (SYNTHGENE)-modified mRNAs were capped using Cap1 Analogue Reagent (TriLink) and further purified by Monarch RNA purification columns (NEB) and resuspended in a TE buffer at the desired concentration. For mRNA encapsulation into LNP, lipid components were dissolved in ethanol at molar ratios of 50:10:38.5:1.5 (ionizable lipid: DSPC: cholesterol: PEG-lipid, all purchased from Firestone Biotechnology). The novel ionizable lipid (YX-02) was designed and patented by Firestone Biotechnologies. The lipid cocktail was mixed with mRNAs dissolved in 10mM citrate buffer (pH4.0) at an N/P ratio of 5.3 :1 and a volume ratio of 3: 1 using a microfluidic-based equipment (INano^TM^L from Micro&Nano Biologics) at a total flow rate of 12 mL/min. Formulations were diluted with PBS and ultrafiltrated using 50-kDa Amicon ultracentrifugal filters. Vaccine formulation was characterized for particle diameter, polymer dispersity index (PDI) and zeta potentials using NanoBrook Omni ZetaPlus (Brookhaven Instruments).

### Protein Expression Assay

Human embryonic kidney (HEK) 293T cells and DC 2.4 cells were cultured in high-glucose Dulbecco’s Modified Eagle Medium (BIOIND, Israel) supplemented with 10% fetal bovine serum (FBS, BIOIND, Israel) and 1% penicillin-streptomycin (NCM Biotech, China). gE-mRNAs were transfected into cells using jetMESSENGER (Polyplus-transfection®) and incubated for 24 hours. Expression of gE protein on cell surface was determined by staining cells with anti-gE mAb (1:200 dilution, Abcam) for 1 hour prior to incubation with PE-anti-human IgG Fc (1:200 dilution, Biolgend) for 30 minutes. Flow cytometric analysis was carried out on Attune NxT (Thermo). Data were analyzed using FlowJo V.10.1 (Tree Star).

### Measurement of gE-specific IgG and IgG subclasses

gE proteins (Acro Biosystems) were coated into 96-well plates (Greiner Bio-One) at a concentration of 50ng/well and incubated overnight at 4C. The plates were washed three times with PBS containing 0.075% Tween-20 (PBST) and blocked for two hours at room temperature (RT) with 2% bovine serum albumin (BSA). Sera samples serially diluted were added and incubated for 2h at RT. For the analysis of murine samples, binding IgG, IgG1, and IgG2c were determined using HRP-conjugated goat-anti-mouse IgG (1:50,000, Abcam), IgG1 (1:5000, Southern Biotech), IgG2c (1:5000, Southern Biotech) Ab for 1 hour at 30C, respectively. For the analysis of NHP sample, binding IgG were determined using HRP-conjugated goat anti-monkey IgG (1:50,000, Abcam) for 1 hour at 30C. TMB substrate was used for development and the absorbance was read at 450 nm wavelength. Endpoint titer was calculated as the dilution factor that emitted an optical density (OD) value above 4.1×background.

### Analysis of memory B cell response

Frequencies of VZV gE-specific class-switched IgD^-^IgM^-^ MBCs were assessed by flow cytometry. For the preparation of gE-probes, biotinylated gE proteins (Acro Biosystems) were conjugated with BV421- or APC-streptavidin (Biolegend) at a molar ratio of 4:1. Cells from mouse spleen or dLN or PBMC from rhesus macaques were first incubated with gE-probes for 20 minutes, and then stained with Fixable Viability Dye eFluorTM 506 (eBioscience) for 5 minutes. After washing, cells were incubated with Fc Receptor blocking reagent (Miltenyi) and antibody cocktails for 20 minutes at 4C in the dark. Flow cytometric analysis was carried out on BD FACSymphony A3 (BD Biosciences). Data were analyzed using FlowJo V.10.1 (Tree Star). A list of antibodies used in this analysis is available in supplemental materials.

### Antigen recall T cell assay

For the evaluation of gE-specific T cell response, a total of 2 million murine splenocytes or rhesus PBMCs were seeded per well into 96-well U-bottom plates and incubated with or without gE protein (2 μg/mL, Acro Biosystems) or gE overlapping peptides (10 μg/mL, 153 peptides, 15mers with 11 aa overlap) in the presence of brefeldin A (BFA, Biolegend) for 8 hours or 16 hours at 37 °C, respectively. Cytokine production of T cells was evaluated by surface and intracellular staining using Fixation/Permeabilization Solution Kit (BD Biosciences). Background cytokine staining was subtracted, as defined by staining in the samples incubated with medium alone. A list of antibodies used in this analysis is available in the Supplemental Table 1 and 2.

### Enzyme-Linked Immunospot (ELISPOT) Assay

Mouse splenocytes or rhesus macaques PBMCs (0.2 million cells per well) were incubated with or without gE protein (1μg/mL, Acro Biosystems) for 20 hours. Frequencies of IFN-γ or IL-2-secreting T cells were assessed using commercial kits (Mabtech) according to the manuals. Spots were developed with BCIP/NBT substrate (Mabtech) and counted using CTL-Immunospot S6 Analyzer. Results were depicted as spot-forming cell (SFC) per million stimulated cells.

### Statistical analysis

Statistical calculations were performed using GraphPad Prism v6.0. Comparisons between groups were determined using two-way analysis of variance (ANOVA) or Mann-Whitney U test. Comparisons between two different time points within same vaccine group were analyzed using 2-tailed paired Wilcoxon’s test. A *p* value less than 0.05 was considered statistically significant (*p ≤ 0.05, **p ≤ 0.01, ***p ≤ 0.001, ****p ≤ 0.0001).

## Results

### Rational design and characterization of a novel VZV mRNA vaccine candidate (ZOSAL)

The glycoprotein E (gE) is a dominant immunogen of VZV and has been widely used as a key antigen candidate for VZV vaccine development. Different from Moderna’s VZV mRNA vaccine encoding for a carboxyl-terminal truncated form of gE antigen [21], we designed our mRNA vaccine encoding the full-length gE antigen considering that efficient T cell epitopes were reported to span the whole protein [26]. Recently, we reported a proprietary artificial intelligence (AI)-based algorithm (LinearDesign) that can design mRNA sequence to achieve optimal folding stability and codon usage that together contribute to high translation efficiency and high vaccine immunogenicity [24]. Using this AI tool, we designed a codon-optimized gE-mRNA sequence based on the wide type sequence from VZV Oka strain (GenBank: AY253715.1). The optimized gE-mRNAs were synthesized using in-vitro transcription (IVT) procedures with N1-methyl-pseudouridine (m1Ψ) modification and showed robust and efficient protein expression upon transfection into HEK-293T and DC2.4 cells (**Figure S1a, b**). VZV mRNA vaccine formulation (named as ZOSAL) was further prepared by encapsulating gE-mRNAs into LNP delivery system using a microfluidics-based procedure. Transmission electron microscopy and dynamic light-scattering analyses confirmed a high homogeneity in both shape and size of the nanoparticle, which demonstrated a particle size of 105.3 ± 1.6 nm with a polymer dispersity index (PDI) below 0.1 analyzed by Dynamic light scattering (**Figure S1c, d**).

### ZOSAL induced robust VZV-specific Ab and memory B cell responses in mice

Immunogenicity of ZOSAL and Shingrix was first evaluated and compared in C57BL/6 mice that were i.m. immunized with escalating doses (1μg, 5μg, 10μg) of ZOSAL or 0.1 human dose of Shingrix on day 0 and day 14 (**Figure 1a**). Following vaccination, we noticed a transient loss of body weight 7 days after booster in mice from the Shingrix group (**Figure S2**), which supported high reactogenicity of Shingrix in line with clinical observation [6,7]. While in contrast, ZOSAL-vaccinated mice showed a steady increase in body weight indicating a good safety profile. With regards to the Ab response, two doses of ZOSAL induced robust levels of gE-specific IgG in a dose-dependent manner. The response was more prominent in mice receiving medium (5μg) or high dose (10μg) mRNA vaccine and was at a comparable level to that induced by Shingrix (**Figure 1b**). Further by analyzing gE-specific IgG subclasses, we found that ZOSAL was more potent at inducing Th1-prone IgG responses that are demonstrated by induction of a lower IgG1 titer and higher IgG2c/IgG1 ratio (**Figure 1c, d**). Fc-mediated Ab functions, including Ab-dependent complement deposition (ADCD) and Ab-dependent neutrophil phagocytosis (ADNP), were also assessed by systems serology approaches using sera collected 14 days after the boost [27]. Despite a distinct IgG subclass composition observed (**Figure 1d**), there was no difference in ADCD (**Figure 1e**) and ADNP (**Figure 1f**) effects between the two vaccine groups. Since Ab function is affected not only by the subclass but also by the patterns of glycosylation of Fc region. Whether ZOSAL and Shingrix induce different glycosylation patterns of Abs awaits further investigation. In addition, we measured the frequencies of class-switched IgD^-^IgM^-^ memory B cells (MBCs) specific to gE antigen in spleens and lymph nodes draining the vaccine injection site (*d*LN) 4 weeks after the boost. Two doses of 5μg or 10μg ZOSAL elicited a significantly higher level of gE^+^ MBCs than Shingrix, which was found in both two lymphoid organs dissected (**Figure 1g, h**). This suggested that ZOSAL was likely to induce a more sustained vaccine response than Shingrix.

**Figure 1.**
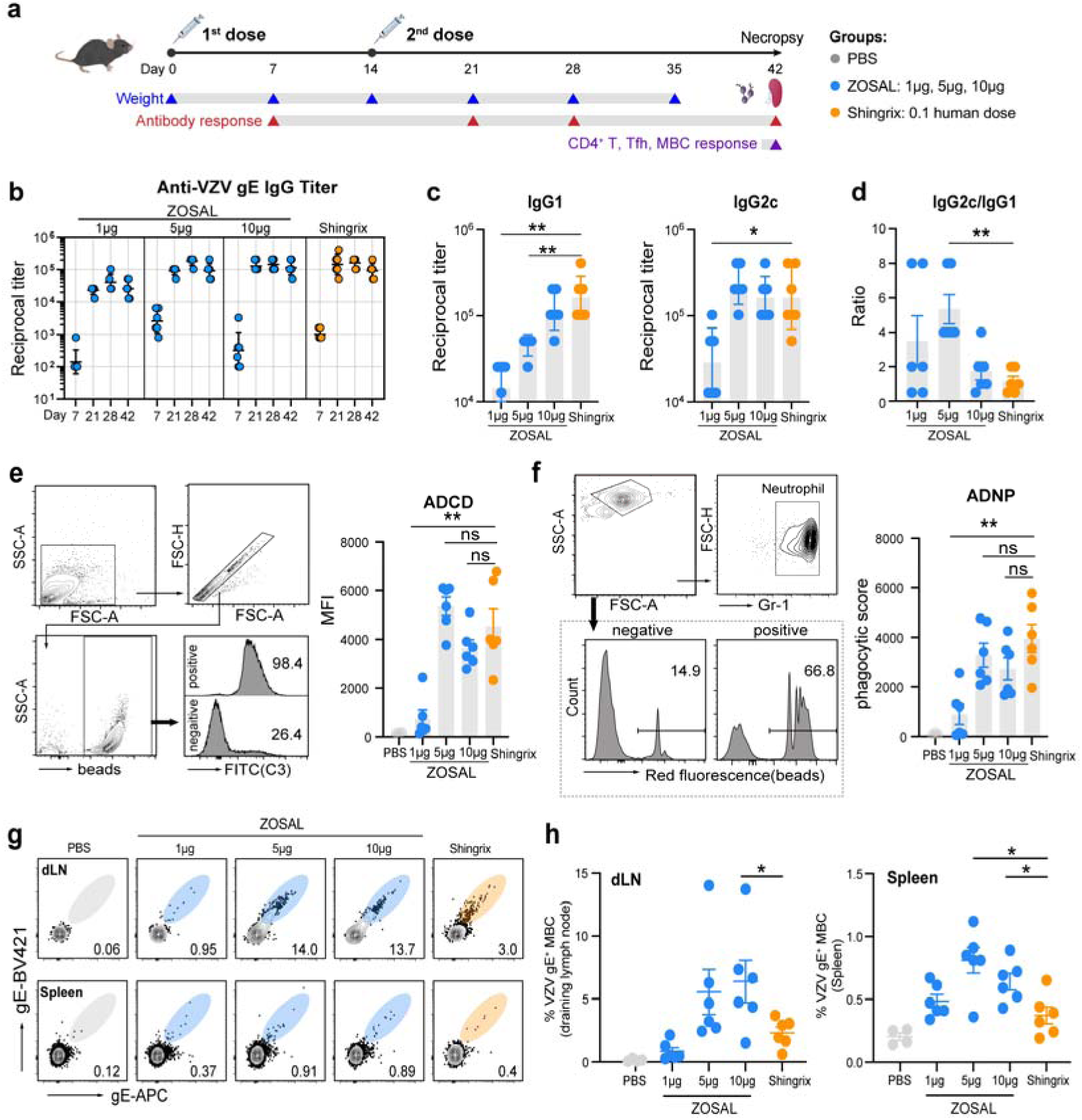
Antibody and memory B cell responses induced by ZOSAL and Shingrix in mice. **a.** Experimental design. C57BL/6 mice (n=6) were i.m. immunized with escalating doses of ZOSAL or 0.1 human dose of Shingrix on day 0 and day 14. Blood draws were taken at the indicated time points. Spleens and dLNs were collected 28 days after the boost. **b.** Anti-gE IgG titers were measured by ELISA and endpoint titers are shown. **c.** Anti-gE IgG1 and IgG2c titers at day 28 were measured by ELISA and endpoint titers are shown. **d.** Ratio of IgG2c/IgG1 is shown. **e-f.** ADCD and ADNP functions of Abs were analyzed using sera collected at day 28. gE-coated microbeads were incubated with diluted and heat-inactivated sera. ADCD (**e)** was detected by fluorescently labeled anti-C3 Abs and MFIs are shown. ADNP (**f**) was determined by beads-positive primary neutrophils and phagocytic scores are shown. **g-h.** Frequencies of class-switched (IgD^-^IgM^-^) gE-specific MBCs in dLNs and spleens were assessed by flow cytometry. Data are shown as mean ± SEM. Mann-Whitney U test was used for statistical analysis. *p ≤ 0.05, **p ≤ 0.01.

### ZOSAL induced stronger gE-specific T cell responses than Shingrix in mice

As Th1-biased CD4^+^ T cell response is critical for the prevention and control of VZV [16,28], we used an intracellular cytokine recall assay to evaluate the induction of gE-specific CD4^+^ T cells and their production of IFN-γ, TNF, IL-2, as well as IL-21 at day 28 after boost immunization. Compared with Shingrix, ZOSAL vaccination elicited significantly higher frequencies of Th1-type CD4^+^ T cells in spleens. Interestingly, the induced T cell response was more prominent in mice receiving low or medium doses of mRNA vaccine (**Figure 2a**), which did not agree with the dose-dependent increasing trend observed in Ab responses described above (**Figure 1b**). Moreover, we detected a stronger induction of gE-specific AIM^+^ (activation-induced marker, OX40 and CD137) CD4^+^ T cells upon antigen stimulation in spleens of ZOSAL-vaccinated mice (**Figure 2b**). Frequencies of cytokine-producing CD8^+^ T cells were also assessed by FACS but showed no clear induction by the two vaccines (data now shown). T follicular helper (Tfh) cells that are specialized in providing helper signals to B cells and critical for germinal center reaction were also measured in parallel in our study. We found that gE-specific Tfh cells defined as OX40^+^CD137^+^CD4^+^CXCR5^+^ T cells were significantly expanded in spleens, especially in mice vaccinated with ZOSAL (**Figure 2c, d**). Moreover, these vaccine-induced Tfh cells showed a clearly activated phenotype indicated by an elevated expression of ICOS (**Figure 2e**). All these together showed that ZOSAL was more potent at inducing VZV-specific T cell responses than Shingrix in mice.

**Figure 2.**
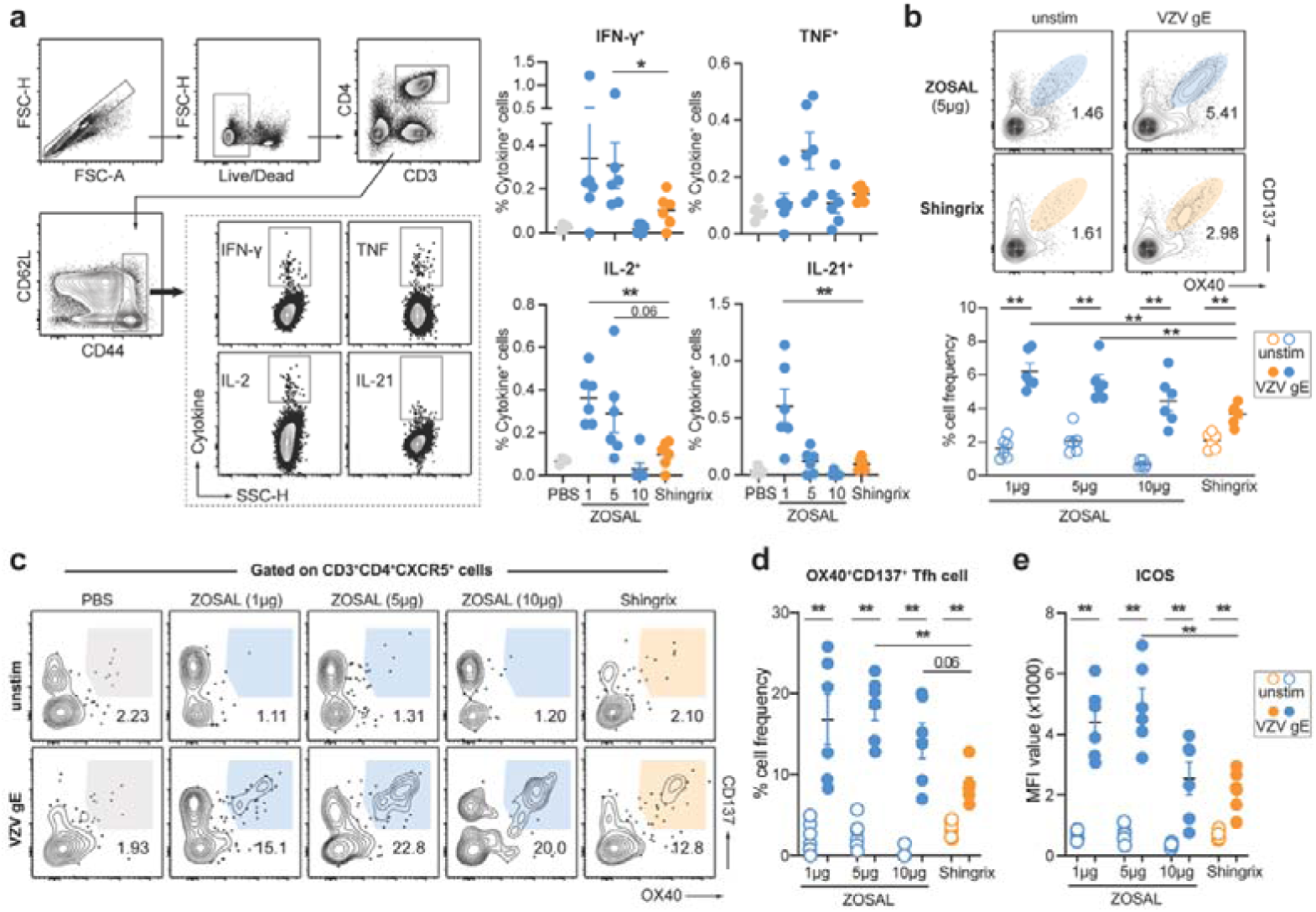
T cell responses induced by ZOSAL and Shingrix in mice. C57BL/6 mice (n=6) were i.m. immunized with escalating doses of ZOSAL or 0.1 human dose of Shingrix on day 0 and day 14. Spleens were collected 28 days after the boost immunization. **a.** Splenocytes were stimulated with or without gE antigen (2 μg/ml) for 8 hours in the presence of Brefeldin A. Frequencies of IFN-γ, IL-2, TNF, or IL-21-secreting CD4^+^ T cells were analyzed by flow cytometry. Data from one representative animal is shown. **b-d.** Splenocytes were stimulated with or without gE antigen (2 μg/ml) for 20 hours. Frequencies of AIM^+^CD4^+^ T cells **(b)** and AIM^+^ Tfh cells (**c-d**) were determined by flow cytometry. **e.** Expression of ICOS on OX40^+^CD137^+^CXCR5^+^CD4^+^ Tfh cells was evaluated. MFI value is shown. Data represent mean ± SEM. Mann-Whitney U test was used for statistical analysis. *p ≤ 0.05, **p ≤ 0.01.

As the incidence for herpes zoster increases with age [8], we also evaluated and compared the immune responses induced by ZOSAL and Shingrix in aged C57BL/6 mice (10-month old) following two doses of vaccination at day 0 and day 14, respectively. While anti-gE IgG titers were at comparable levels between two vaccine groups (**Figure S3a**), aged mice immunized with ZOSAL demonstrated a lower level of IgG1, a higher level of IgG2c, and a remarkably higher IgG2c/IgG1 ratio (**Figure S3b, c**). Regarding gE-specific T cell responses, ZOSAL induced a significantly higher level of IFN-γ or IL-2-producing T cells than Shingrix (**Figure S3d, e**). These data were in line with the above findings in adult mice and further supported the superior immunogenicity of ZOSAL.

### Innate immune responses induced by ZOSAL and Shingrix in mice and rhesus macaques

Sufficient stimulation of innate immunity by vaccination is fundamental to the generation of high-quality vaccine-specific adaptive responses. To this end, we next studied the innate immune responses in C57BL/6 mice that were i.m. administered with different doses of ZOSAL (1μg, 5μg, 10μg) or corresponding equivalent amount of empty LNP, which were benchmarked to Shingrix. We found that ZOSAL induced a significant dose-dependent activation of two major dendritic cell (DC) subsets, cDC1 and cDC2 both in the spleens and dLNs at early time point (12 hours) after vaccine administration (Figure S4a-b). Shingrix induced strong activation of DCs in the dLNs but not in the spleens (Figure S4b). In addition, empty LNP vector also induced splenic cDC1 and cDC2 activation, albeit at a much lower level than LNP containing mRNA (ZOSAL), which suggested an immune-stimulatory effect of LNP as previously reported [29,30]. Serum alanine aminotransferase (ALT) and aspartate aminotransferase (AST) levels were also assessed, which were not altered early after vaccination and therefore suggested there was no acute liver toxicity induced (Figure S4c).

Nonhuman primate serves as an important animal model for preclinical testing of vaccines due to their genetic and physiological similarities to humans. To better assess our mRNA vaccine and gain mechanistic insights into the generation of VZV immunity by two different vaccine modalities, we next performed an in-depth characterization of the vaccine responses induced by ZOSAL and Shingrix in rhesus macaques (**Figure 3a**). Two doses of ZOSAL (100 μg) or Shingrix (human dose) were i.m. administered at an interval of 4 weeks. We first measured the early innate immune responses after vaccine administration by monitoring the fluctuation of distinct leukocyte subsets. 24 hours after prime immunization, both ZOSAL and Shingrix were shown to induce a rapid and transient decrease in circulating lymphocytes, accompanied by a transient increase in neutrophils and monocytes (**Figure S5**). Among the expanded monocytes, there was a noticeable increase of CD14^+^CD16^+^ intermediate monocytes not only in cell frequency but as a proportion within the monocyte compartment (**Figure 3b, c**), which was slightly more pronounced in ZOSAL vaccine group and was found both after prime and booster dose. These findings coincided with what we have previously reported for several other vaccines including a COVID-19 mRNA vaccine [31], a rabies mRNA vaccine [32] and a protein-based subunit malaria vaccine [33], which reflect a strong innate immune activation.

**Figure 3.**
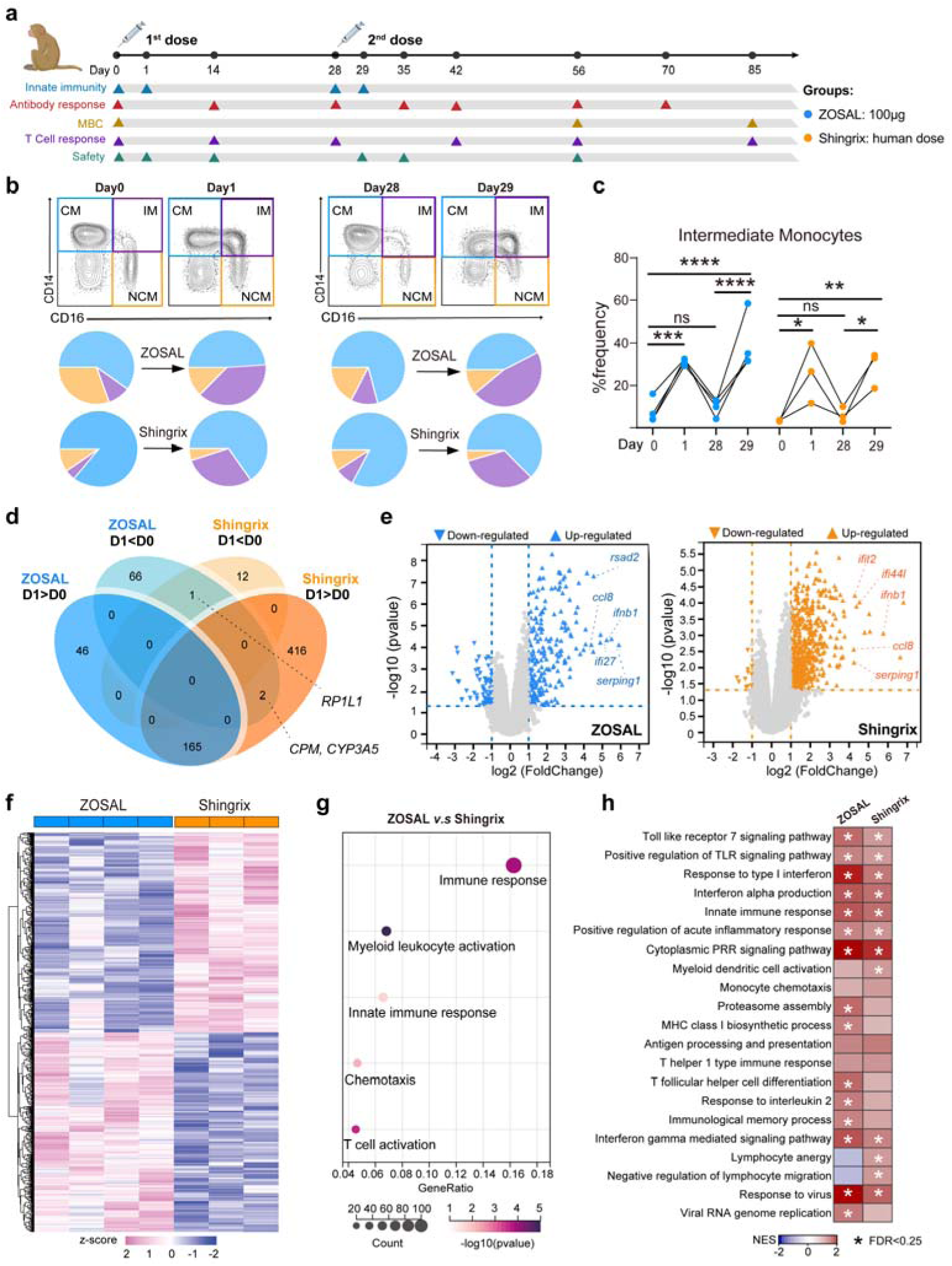
Alteration of intermediate monocytes and modulation of gene expression after ZOSAL and Shingrix vaccination in rhesus macaques. **a.** Experimental design. Rhesus macaques were i.m. immunized twice with ZOSAL (n=4) or Shingrix (n=3) at an interval of 4 weeks. Blood and serum samples were taken at the indicated time points for analysis. **b.** Differentiation of three monocyte subsets 24 hours after prime and boost immunization is shown. Classical monocyte (CM), intermediate monocyte (IM), or nonclassical monocyte (NCM). Pie charts indicate percentage of each monocyte subset out of total monocytes. **c.** Frequencies of CD14^+^CD16^+^ intermediate monocytes are shown. **d-h.** Transcriptomic analyses of PBMCs isolated 24 hours post the boost immunization. **d.** Venn diagram indicates the number of altered genes shared by the two vaccine groups. D1CD0 and D1CD0 represent up-regulated and down-regulated genes, respectively. **e.** Volcano plots display genes that were altered by ZOSAL or Shingrix vaccination. Criteria used are *p* value < 0.05 calculated using a paired two-tailed Student’s *t* test and at least 2-folded change after vaccination. Representative genes are annotated. **f.** Heatmaps of significantly altered DEGs by ZOSAL or Shingrix are shown. Values in heatmaps are z-score standardized. **g**. DEGs in ZOSAL-vaccinated animals versus Shingrix-vaccinated animals were employed for GO analysis, with focus on the indicated GO biological process terms. **h**. GSEA analysis using all DEGs. Each box represents a specific module and colors indicate normalized enrichment score (NES). Asterisk denoted in the box represents False Discovery Rate (FDR) values < 0.25. Two-way ANOVA was used for statistical analysis in figure c. *p ≤ 0.05, **p ≤ 0.01, ***p ≤ 0.001, ****p ≤ 0.0001.

To gain a more in-depth understanding of the innate immune activation, we performed transcriptomic analyses on PBMCs isolated 24 hours post the boost immunization. Profiles and changes in gene expression were thoroughly evaluated using bioinformatic approaches. After vaccination, there was a considerable overlap of 165 genes showing ≥ 2-fold increase and only one gene (*RP1L1*) showing ≤ 0.5-fold decrease that were shared by the two vaccine groups when compared with matched pre-vaccination (day 0) samples (**Figure 3d**). Two other genes, namely *CPM* and *CYP3A5* were down-regulated after ZOSAL vaccination but in contrast were increased upon Shingrix vaccination. The exact physiological outcomes of changes in *CYP3A5* expression by two different vaccines remains unknow, but it was likely to be associated with drug metabolization and the resulting toxicity since *CYP3A5*-encoded liver enzyme is critical to metabolize and detoxify xenobiotics [34,35]. Genes that were altered in their expression by the two vaccines were further identified, amongst which a large number of genes (*ccl8, ifnb1, serping 1*, etc) related to interferon (IFN) response, complement pathway and chemotaxis were up-regulated and commonly shared by the two groups (**Figure 3e**). And it was clearly observed that ZOSAL and Shingrix showed distinct impact on gene expression patterns (**Figure 3f**). Further, differentially expressed genes (DEGs) were subjected to a Gene Ontology (GO) analysis using GO biological process database and were found to be highly enriched in pathways related to “immune response”, “myeloid leukocyte activation”, “innate immune response”, “chemotaxis” and “T cell activation” (Figure 3g). These DEGs were depicted in the Supplemental Table 3. Furthermore, Gene Set Enrichment Analysis (GSEA) was performed for in-detail characterization of their functions and enrichments in gene modules. Modules including TLR signaling, Type-I IFN response, cell chemotaxis, antigen processing and presentation, lymphocyte function, etc that are closely related to the generation of vaccine responses were in general highly upregulated in both two groups (**Figure 3h, Table S4**), which was more prominent in ZOSAL-vaccinated rhesus macaques. Interestingly, although Shingrix is potent at inducing strong T cell responses, genes associated with “lymphocyte anergy” and “negative regulation of lymphocyte migration” concomitantly increased. While in contrast, these two modules were clearly decreased upon ZOSAL vaccination (**Figure 3h**). This further suggested the superiority of ZOSAL over Shingrix in eliciting stronger T cell immunity in rhesus macaques as we studied later (**Figure 5**).

### ZOSAL induced comparable levels of Ab and MBC responses to Shingrix in rhesus macaques

We next evaluated the VZV-specific Ab and B cell responses induced by ZOSAL and Shingrix in rhesus macaques. Consistent with what we have found earlier in mouse model (**Figure 1**), both two vaccines were able to induce detectable levels of anti-gE IgG after prime immunization, and the titers increased remarkably reaching a comparable and peak level 7 days after the boost (**Figure 4a**). Class-switched gE-specific MBCs in PBMCs were also analyzed according to the indicated gating strategy (**Figure S6a**), which were robustly induced by both two vaccines at an equivalent level and their cell frequencies correlated well with Ab titers (**Figure 4b, c**). Functions of Abs including Ab-dependent cellular phagocytosis (ADCP) by THP-1 cells and ADCD effects were also evaluated using sera collected at different time points after vaccination. No clear difference was observed regarding the functions of Abs induced by ZOSAL and Shingrix, which was in line with our findings in C57BL/6 mice (**Figure 1e, f**).

**Figure 4.**
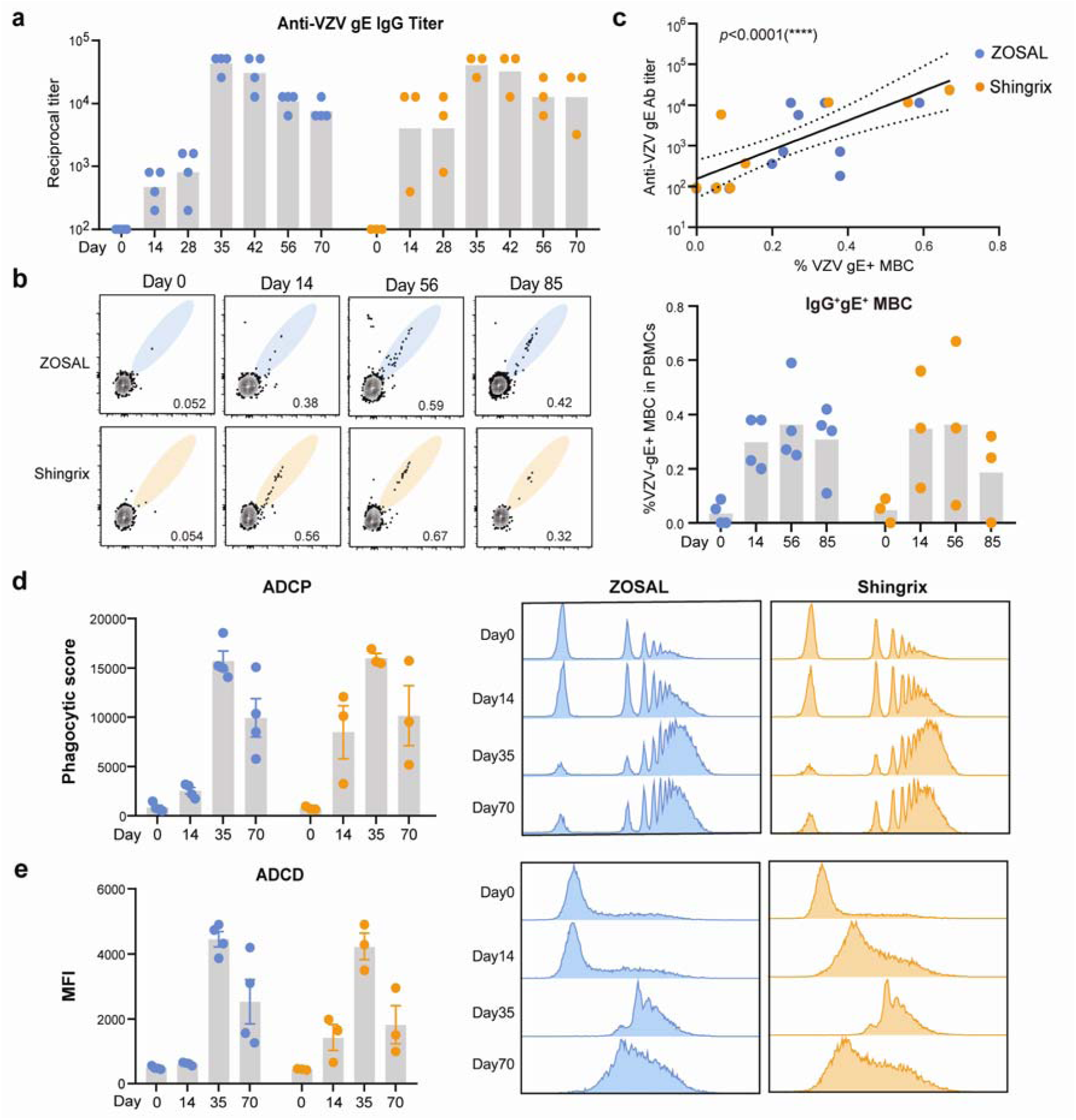
Antibody and memory B cell responses induced by ZOSAL and Shingrix in rhesus macaques. Rhesus macaques were i.m. immunized twice with ZOSAL (n=4) or Shingrix (n=3) at an interval of 4 weeks. **a.** Endpoint titers of anti-gE IgG were measured by ELISA longitudinally. **b.** Frequencies of class-switched IgD^-^IgM^-^ gE-specific MBCs in PBMCs were assessed by flow cytometry. Data from representative animals (left panel) and cell frequencies are shown as mean ± SEM (right panel). **c**. Correlation of class-switched gE^+^ MBCs and anti-gE IgG titers. **d.** gE-coated microbeads were incubated with diluted and heat-inactivated sera, followed by incubation with THP-1 cells. ADCP effect of Abs was determined as frequencies of beads-positive cells and phagocytic scores are shown. **e**. ADCD effect of Abs was detected by fluorescently labeled anti-C3 Abs and MFIs are shown. Pearson’s correlation analysis was used. Data are shown as mean ± SEM. ****p ≤ 0.0001.

### ZOSAL was more potent at inducing a Th1-biased CD4^+^ T cell response than Shingrix in rhesus macaques

Given that Th1-type T cells, not Abs, mediate the protection against VZV [10–12], we gave more attention to the T cell responses elicited. Using ELISpot assays, we found that a single prime dose of ZOSAL could induce a moderate level of IFN-γ or IL-2 secreting T cells at a higher magnitude than Shingrix (**Figure 5a, b**). Such response was markedly enhanced following the boost immunization, especially in animals receiving ZOSAL vaccination. As previous study has validated a more critical role of CD4^+^ T cells rather than CD8^+^ T cells in protection against varicella virus [14,16], we further used a well-established intracellular cytokine recall assay (**Figure S6b)** to evaluate the induction of gE-specific CD4^+^ T cells producing Th1-type cytokines (IFN-γ, TNF, IL-2) upon stimulation by gE overlapping peptides pool. Two or four weeks after the boost immunization, ZOSAL vaccination elicited significantly higher frequencies of Th1-type CD4^+^ T cells than Shingrix, although no statistical significance was observed due to limited number of animals used. Frequencies of cytokine-producing CD8^+^ T cells were also analyzed in parallel, which showed no clear induction in both two vaccine groups (**Figure S7**). Combined with the findings in mouse model (**Figure 2**), these results collectively showed that ZOSAL was superior at inducing VZV-specific CD4^+^ T cell immunity than Shingrix.

**Figure 5.**
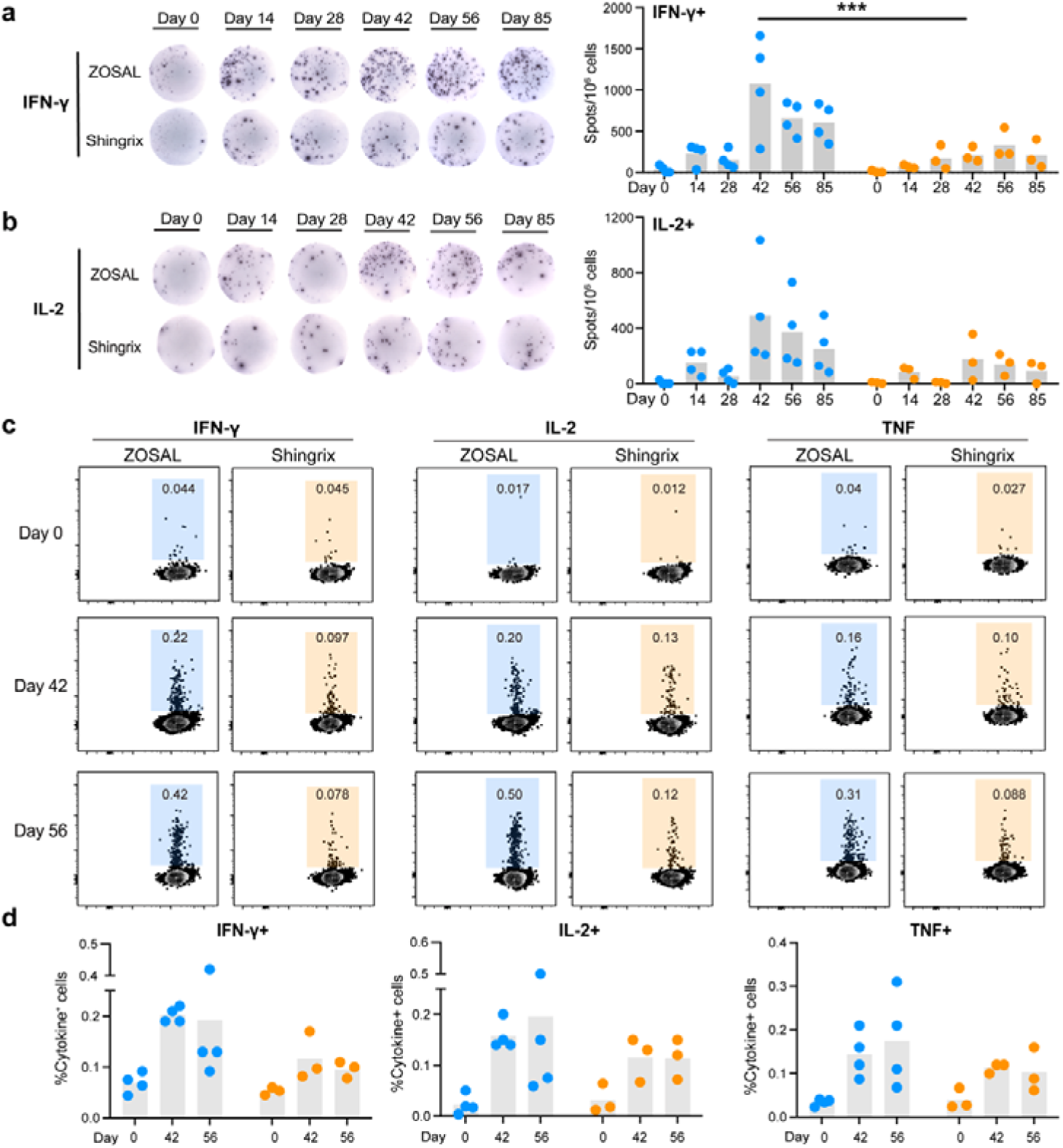
A stronger Th1-biased CD4^+^ T cell response was induced by ZOSAL than Shingrix in rhesus macaques. Rhesus macaques were i.m. immunized twice with ZOSAL (n=4) or Shingrix (n=3) at an interval of 4 weeks. **a-b.** PBMCs collected at different time points before and after vaccination were stimulated with gE antigen (2 μg/ml) for 20 hours. Frequencies of IFN-γ or IL-2-secreting T cells at the indicated time points were measured by ELISpot. Data from representative animals are shown. Numbers of gE-specific cytokine-producing T cells were enumerated and are shown as spots per million stimulated cells**. c.** PBMCs were stimulated with or without gE overlapping peptides pool (10 μg/ml) for 16 hours in the presence of Brefeldin A. Frequencies of IFN-γ, IL-2, TNF-secreting CD4^+^ T cells were analyzed by flow cytometry. Data from representative animals are shown. **d.** Quantification of cytokine-producing CD4^+^ T cells upon antigen stimulation. Two-way ANOVA with multiple comparisons tests was used for analysis of statistical significance. ***p ≤ 0.001.

### Correlation matrix and cluster analysis of immune parameters identified

Longitudinal sample collection and the above comprehensive analyses gave us an opportunity to elucidate the correlations among key immune parameters identified, which help us better understand the generation of vaccine-induced VZV immunity. We therefore took the step further to perform a multivariate nonparametric correlation analysis using 44 variables measured in our study followed by hierarchical clustering as described previously [33]. Early innate immune parameters correlating with adaptive vaccine responses were particularly emphasized. Three clusters were identified in our analysis (**Figure 6a**). Amongst them, cluster 1 that contains genes relevant to IFN response (*mx1, isg20*), antigen presentation (*b2m, cd80*), and TLR7 signaling pathway showed highly positive correlations with gE-specific Th1-type T cell responses. Interestingly, cluster 3 that includes neutrophil count and neutrophil/lymphocyte ratio measured at day 1, ADCP function of Abs at day 35, etc showed negative correlations with cluster 1 representing T cell responses. The most critical endpoints in our study, that is frequencies of Th1-type T cells, were further isolated for a better visualization of their associating variables. It was clearly demonstrated that vaccine-induced T cell responses were highly and positively correlated with the expression of IFN-stimulated genes. While genes related to chemotaxis (ccr6, ccr9), ADCP function of Abs, and neutrophil/lymphocyte ratio were the major negative regulating factors (**Figure 6b)**. Altogether, we identified several early immune factors that can predict the magnitude of T cell response and potentially predict vaccine efficacy.

**Figure 6.**
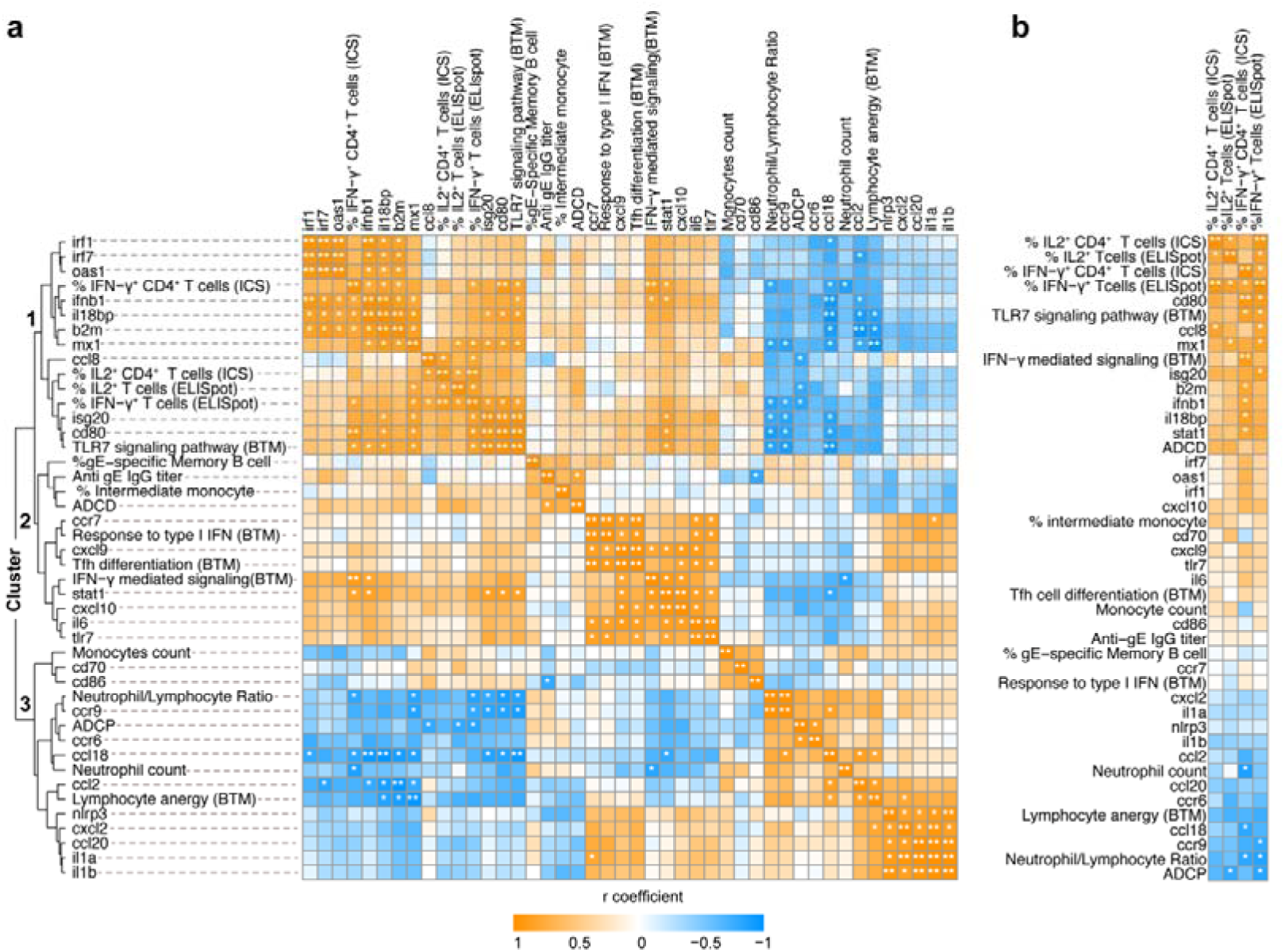
Correlation and cluster analysis of immune parameters. **a.** A multivariate nonparametric Spearman’s test was used to analyze the correlation among 44 immune parameters measured in the study. Heatmap shows the correlation coefficient with four clusters denoted. *p ≤ 0.05, **p ≤ 0.01. **b.** Immune parameters correlating with gE-specific Th1 T cell responses were isolated and shown.

### ZOSAL demonstrated a superior safety profile than Shingrix in rhesus macaques

A promising vaccine candidate should possess a high safety profile. Despite the fact that Shingrix is highly effective to prevent zoster, clinical trials and post-marketing surveillance have reported a high local and systemic reactogenicity of Shingrix [36]. In this study, we preliminarily evaluated safety properties of ZOSAL by monitoring levels of various serum biochemical parameters following vaccination, including liver and kidney enzymes. In general, both ZOSAL and Shingrix vaccinated macaques were well-tolerated and these parameters remained stable after vaccination (**Figure 7a**). Further, by revisiting the GSEA analyses on RNA-sequencing data, we noticed that modules associated with platelet degranulation, aggregation, activation, and pain responses were largely increased upon Shingrix vaccination, which in contrast were unchanged or even decreased after ZOSAL vaccination. Several genes that are typically relevant to platelet activation and pain response were further isolated and shown (**Figure 7b, c**). Collectively, our data indicated a high-level safety property of ZOSAL that is even superior over licensed vaccine.

**Figure 7.**
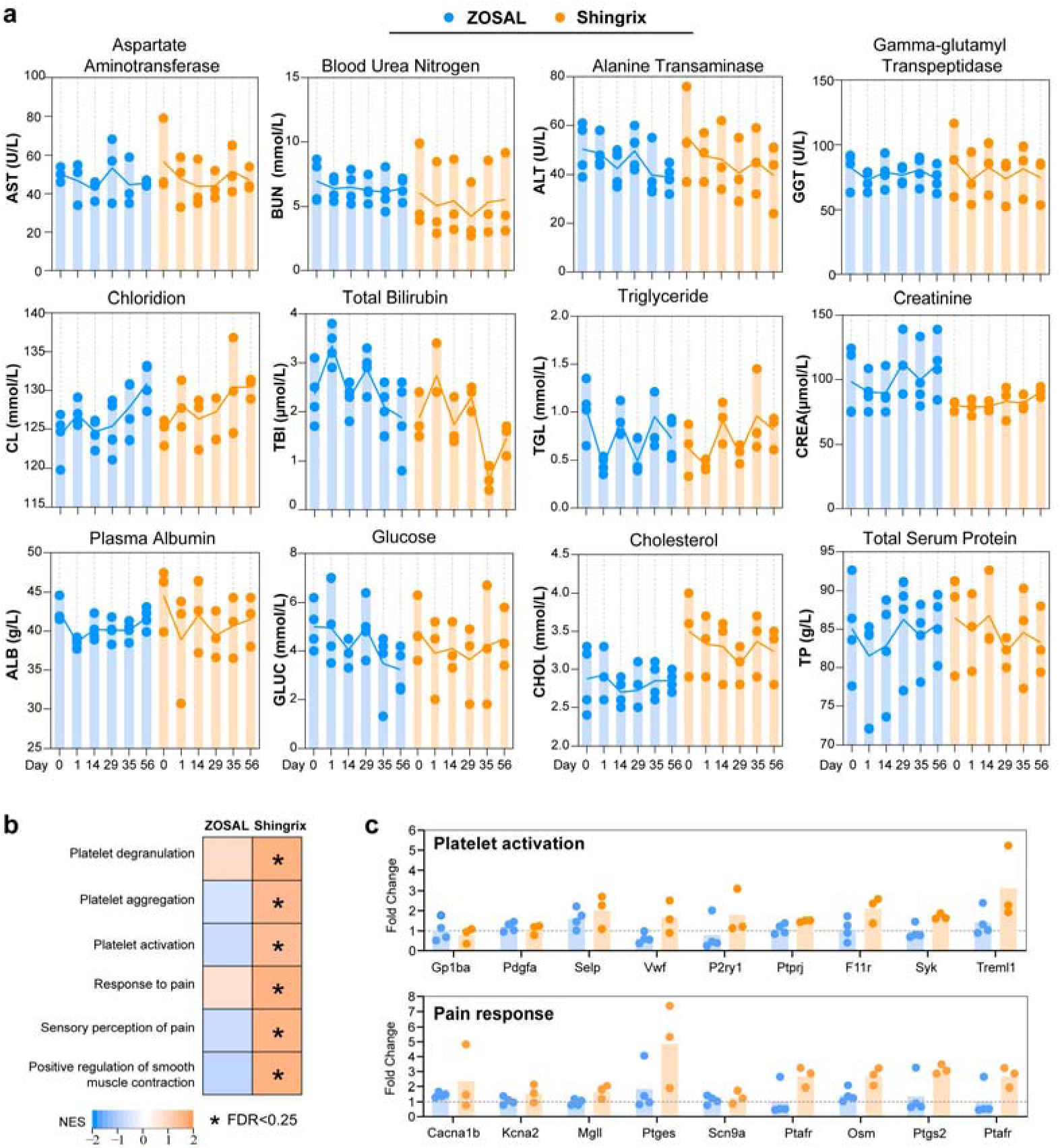
A preliminary evaluation of safety profiles of ZOSAL and Shingrix in rhesus macaques. Rhesus macaques were i.m. immunized twice with ZOSAL (n=4) or Shingrix (n=3) at an interval of 4 weeks. **a.** Levels of serum biochemical parameters at the indicated time points were measured. **b.** GSEA analysis revealed distinct differences in the level of indicated modules related to platelet function and pain response. Each box represents a specific module and colors indicate normalized enrichment score (NES). Asterisk denoted in the box represents False Discovery Rate (FDR) values < 0.25. **c.** Expression of indicated genes after vaccination. Data is shown as fold change normalized to pre-vaccination.

## Discussion

Herpes zoster remains an important global health issue given the continuously rising incidence and limited licensed vaccines available [9]. Shingrix, an adjuvanted recombinant subunit vaccine, has a remarkably high protective efficacy and therefore has been dominating the market since its licensure. However, high reactogenicity that is likely caused by the strong AS01B adjuvant component remains to be a major concern on Shingrix [37,38]. There is therefore a critical need for novel or improved VZV vaccines with high efficacy and low reactogenicity.

VZV-specific T cell response is solely critical for controlling latent reactivation and therefore must be considered in next-generation rational VZV vaccine development [12,13]. By now it is widely-accepted that mRNA vaccines are potent at inducing T cell responses. Both CD4^+^ and CD8^+^ T cell responses have been shown to be induced at higher levels by mRNA vaccines than induced by other protein vaccine platforms [39]. Therefore, for infections like VZV, mRNA platform may be ideal to be employed for vaccine development. In the present study, we developed a novel VZV mRNA vaccine candidate (ZOSAL) and performed multiple assays to thoroughly evaluate vaccine immunogenicity and safety aspects with a side-by-side comparison with Shingrix. In adult mice, aged mice and rhesus macaques, ZOSAL demonstrated superior immunogenicity and safety in multiple aspects over Shingrix, especially in the induction of strong T cell immunity. Moreover, we took the step further to obtain mechanistic insights into the generation of vaccine-induced T cell response and identified the associating immune correlates.

It has been reported that host immune responses to VZV infection involves both Ab and T cell responses [12,40]. While VZV-specific T cell responses decline with age and their frequencies inversely correlate with the incidence or disease severity of HZ, the anti-VZV Abs largely remained stable or even increased with age and showed no association with HZ incidence or severity [41,42]. Moreover, increased risk of HZ in immunocompromised individuals was associated with impaired T cell immunity but not with anti-VZV IgG titers [43]. Although these indicated that VZV-specific Abs are dispensable in protection against VZV, we still measured the Ab and B cell responses in our study, which we believe are critical indicators reflecting the phenotype and longevity of vaccine response. We found that ZOSAL elicited a comparable level of gE-specific IgG to Shingrix in mice, and the induced IgG subclass was much skewed towards a Th1-type IgG2c which suggested a more Th1-prone vaccine response induced by ZOSAL. Despite the distinct IgG subclass composition induced by the two vaccines, no clear difference in the effector functions of Abs were demonstrated. Previous studies have reported that Ab function is affected not only by the subclass but also by the patterns of glycosylation of Fc region [44–46], and distinct vaccine regimens induce different IgG glycosylation profiles [47]. It should be noted that ZOSAL and Shingrix represent two types of vaccines and their interactions with immune system and the resulting immune activation profiles are quite different. Whether ZOSAL and Shingrix induce different glycosylation patterns of Abs and how this influences vaccine efficacy is an interesting topic to be further investigated. Neutralizing capacity of Abs were not assessed in our study due to technical reason, and the exact role of neutralizing Abs in the control of VZV infection remains yet unclear. Regarding the B cell response, class-switched gE^+^ MBCs induced by ZOSAL was at a significantly higher level than that induced by Shingrix, which suggested ZOSAL was likely to induce a more sustained vaccine response than Shingrix, although not directly detected in our study.

T cell immunity was particularly focused in our study due to the aforementioned reason. In different animal models tested, ZOSAL demonstrated a significantly higher potency at eliciting VZV-specific Th1-type CD4^+^ T cell responses than Shingrix. However, we did not observe a clear induction of VZV-specific CD8^+^ T cells by the two vaccines. This was possibly due to the low frequency of VZV gE-specific CD8^+^ T cells or the low sensitivity of immune assays used. Moreover, gE (ORF68) antigen may contain limited CD8^+^ T cell epitopes since it has been reported that VZV-specific CD8^+^ T cells were largely reactive to ORF9, not to other VZV antigens including gE and gB [48]. Another aspect that should be noted is that the VZV-specific T cells induced by ZOSAL did not show a dose-dependent pattern, which was in contrast to the induction of Abs. Similar findings have also been reported previously in many other mRNA vaccine studies [49–51]. So far, mechanisms of action of mRNA vaccines remain largely unclear and await further in-depth investigation.

Effective generation of pathogen-specific T cells is largely determined by sufficient innate immune stimulation, especially the events associated with antigen presentation. Differences in the early innate responses induced by the two vaccines were in-depth characterized using nonhuman primate model. Both ZOSAL and Shingrix exhibit powerful abilities to activate innate immune compartments, in particular Type-I IFN signaling and antigen processing/presentation. Although we did not observe a striking difference between the two vaccine groups, this was quite expected since the adjuvant component in Shingrix that contains liposomes formulated with cholesterol, monophosphoryl lipid A (MPL), and QS21 were highly immune-stimulatory to favor the generation of Th1-type vaccine responses [52]. However, it should always be aware that innate activation is a “double-edged sword” and has to be well-controlled to an appropriate degree to aid in the generation of high-quality adaptive vaccine responses and at the same time ensure the safety. Over-activated type-I IFN response has also been reported to be deleterious to the generation of Tfh response favoring Ab production [53]. While, this may unlikely occur to our mRNA vaccine since ZOSAL demonstrated superior ability in eliciting both class-switched gE^+^ MBCs and Tfh cells than Shingrix in mice.

In this study, we also performed a correlation analysis of 44 immune parameters to better understand the generation of vaccine-induced VZV immunity. It is easy to understand the strong positive correlations between expression of IFN-relevant genes and the magnitude of T cell responses. While interestingly, a few parameters including neutrophil/lymphocyte ratio at 24h after vaccination, ADCP function of Abs, etc were negatively associated with T cell responses. Previously, we have reported that number of circulating neutrophils increased rapidly and transiently 24 hours after influenza or rabies mRNA vaccination in rhesus macaques [32,54]. Neutrophils are potent at phagocytosing LNP-mRNA particle but have a very limited capability to translate mRNA into the protein antigens [54]. This may cause an ineffective usage of injected mRNA vaccine that negatively affect the generation of vaccine responses.

We also took advantage of the rhesus macaque study to preliminarily assess safety aspects of our VZV mRNA vaccine. A set of genes associated with platelet activation and pain responses were upregulated after Shingrix vaccination, but not by ZOSAL. Considering that aged individuals are main target population of zoster vaccine and these cohorts are more susceptible to cardiovascular diseases, the inappropriate platelet activation induced by Shingrix may pose a potential risk. Besides, the increased expression of pain-related genes also supported the clinical findings that local injection site pain is the most common solicited adverse event in Shingrix recipients [36]. In addition, it should be noted that our mRNA vaccine encodes full-length gE, which is in contrast with Shingrix that contains truncated form of gE (1-544 aa) [55]. The rationale behind this design was mainly due to the fact that C-terminal domain of gE also contains efficient T cell epitopes that contributes to a broader induction of VZV-specific T cells [26]. However, further investigation would be necessary to address whether using full-length gE as vaccine immunogen poses potential safety risks.

There are also some weaknesses of our study that remain to be further addressed. Dose titration of ZOSAL in rhesus macaques was not performed which was largely due to the limited source of animals and extraordinary high cost. However, this is critical to guide the further studies on ZOSAL using a more optimal dose especially in the potential clinical studies. In addition, we were not able to perform viral challenge study to directly assess the protective efficacy of ZOSAL, although the superior T cell immunity induced by ZOSAL may suggest a superior protection. In fact, there are no adequate animal models of VZV infection that can fully recapitulate VZV reactivation in humans due to the strict host-specificity of VZV infection [56].

To summarize, we have developed a novel VZV mRNA vaccine candidate that has superior immunogenicity and safety property over licensed vaccine in mice and rhesus macaques. To our knowledge, this is the first study that provides high-resolution profiling and comparison of a VZV mRNA vaccine (ZOSAL) and Shingrix side-by-side in both mice and rhesus macaques. Importantly, our data generated from nonhuman primate model that is much appreciated for its high translational value for human vaccine responses will warrant more investigations of the mRNA platform in the development of next-generation herpes zoster vaccines.

## Data availability

All data are available upon reasonable request to the corresponding authors.

## Acknowledgements

This work was supported by the Natural Science Foundation of Jiangsu Province (BK20221031 to A.L.), the National Science Foundation of China (32200764, to A.L; 82001687, to J.H.Z.), the Fundamental Research Funds for the Central Universities (2632022YC01, to A.L.), Shandong Provincial Natural Science Foundation for The Excellent Youth Scholars (No. ZR2023YQ066 to H.Z.), the Open Project of State Key Laboratory of Natural Medicines (No. SKLNMKF202309 to H.Z), and the State Key Laboratory of Microbial Technology Open Projects Fund (No. M2023-13 to H.Z).We thank Min-Hui Sun, Qin-Sheng Dai and Jia Li for technical support from Target Discovery Center of China Pharmaceutical University, and also thank the Core Facility of Shandong University for instrumental and technical support on this study. The authors also thank Firestone Biotechnology (Shanghai) for technical assistance in the development of novel ionizable lipids and thank Baidu, Inc. for authorizing Dr. Lin’s lab to use LinearDesign Algorithm freely for commercial and non-commercial purpose in this study. We thank Dr. Liang Zhang for running LinearDesign Algorithm (Baidu) to codon-optimize the mRNA sequence.

## Author contributions

A.L. and Y.Y. designed the project; L.H., T.Z., L.B., W.Z., H.Z., Y.G., X.Z., C.W., Z.D., S.S., W.M., Z.L., L.S., J.H. and A.L. performed the experiments and analysis; H.Z., L.M., Q.D., J.L., G.W., L.W., S.F. and K.L. provided methodological and technical support; A.L., L.H., Y.Y., K.L. discussed the data. L.H., T.Z., L.B., G.W., A.L. performed the revision; A.L., L.H. and T.Z. wrote the manuscript. All authors have read and approved the manuscript.

## Conflict of interest

J.H. is an employee of Firestone Biotechnologies, which focuses on the development of novel ionizable lipids and LNP formulation. Firestone has filed a patent on the novel ionizable lipid (YX-02) used in this study. All other authors declare no conflict of interest.

## Notes

### Summary of Updates

We have updated some of the figures in this revised version.

## References

[1] Zerboni L, Sen N, Oliver SL, et al. Molecular mechanisms of varicella zoster virus pathogenesis. Nat Rev Microbiol. 2014 Mar; 12(3):197–210.

[2] Nordén R, Nilsson J, Samuelsson E, et al. Recombinant Glycoprotein E of Varicella Zoster Virus Contains Glycan-Peptide Motifs That Modulate B Cell Epitopes into Discrete Immunological Signatures. Int J Mol Sci. 2019 Feb 22; 20(4).

[3] Chen T, Sun J, Zhang S, et al. Truncated glycoprotein E of varicella-zoster virus is an ideal immunogen for Escherichia coli-based vaccine design. Sci China Life Sci. 2023 Apr; 66(4):743–753.

[4] Lee SJ, Park HJ, Ko HL, et al. Evaluation of glycoprotein E subunit and live attenuated varicella-zoster virus vaccines formulated with a single-strand RNA-based adjuvant. Immun Inflamm Dis. 2020 Jun; 8(2):216–227.

[5] Wang L, Zhu L, Zhu H. Efficacy of varicella (VZV) vaccination: an update for the clinician. Ther Adv Vaccines. 2016 Jan; 4(1-2):20–31.

[6] Lal H, Cunningham AL, Godeaux O, et al. Efficacy of an adjuvanted herpes zoster subunit vaccine in older adults. N Engl J Med. 2015 May 28; 372(22):2087–96.

[7] Cunningham AL, Lal H, Kovac M, et al. Efficacy of the Herpes Zoster Subunit Vaccine in Adults 70 Years of Age or Older. N Engl J Med. 2016 Sep 15; 375(11):1019–32.

[8] Yawn BP, Gilden D. The global epidemiology of herpes zoster. Neurology. 2013 Sep 3; 81(10):928–30.

[9] Harpaz R, Ortega-Sanchez IR, Seward JF. Prevention of herpes zoster: recommendations of the Advisory Committee on Immunization Practices (ACIP). MMWR Recomm Rep. 2008 Jun 6; 57(Rr-5):1–30; quiz CE2-4.

[10] Asada H. VZV-specific cell-mediated immunity, but not humoral immunity, correlates inversely with the incidence of herpes zoster and the severity of skin symptoms and zoster-associated pain: The SHEZ study. Vaccine. 2019 Oct 16; 37(44):6776–6781.

[11] Steain M, Sutherland JP, Rodriguez M, et al. Analysis of T cell responses during active varicella-zoster virus reactivation in human ganglia. J Virol. 2014 Mar; 88(5): 2704–16.

[12] Weinberg A, Levin MJ. VZV T cell-mediated immunity. Curr Top Microbiol Immunol. 2010; 342:341–57.

[13] Weinberg A, Zhang JH, Oxman MN, et al. Varicella-zoster virus-specific immune responses to herpes zoster in elderly participants in a trial of a clinically effective zoster vaccine. J Infect Dis. 2009 Oct 1; 200(7):1068–77.

[14] Haberthur K, Engelmann F, Park B, et al. CD4 T cell immunity is critical for the control of simian varicella virus infection in a nonhuman primate model of VZV infection. PLoS Pathog. 2011 Nov; 7(11): e1002367.

[15] Traina-Dorge V, Palmer BE, Coleman C, et al. Reactivation of Simian Varicella Virus in Rhesus Macaques after CD4 T Cell Depletion. J Virol. 2019 Feb 1; 93(3).

[16] Park HB, Kim KC, Park JH, et al. Association of reduced CD4 T cell responses specific to varicella zoster virus with high incidence of herpes zoster in patients with systemic lupus erythematosus. J Rheumatol. 2004 Nov; 31(11):2151–5.

[17] Pardi N, Hogan MJ, Porter FW, et al. mRNA vaccines - a new era in vaccinology. Nat Rev Drug Discov. 2018 Apr; 17(4):261–279.

[18] Verbeke R, Hogan MJ, Loré K, et al. Innate immune mechanisms of mRNA vaccines. Immunity. 2022 Nov 8; 55(11):1993–2005.

[19] Sahin U, Karikó K, Türeci Ö. mRNA-based therapeutics--developing a new class of drugs. Nat Rev Drug Discov. 2014 Oct; 13(10):759–80.

[20] Li C, Lee A, Grigoryan L, et al. Mechanisms of innate and adaptive immunity to the Pfizer-BioNTech BNT162b2 vaccine. Nat Immunol. 2022 Apr; 23(4):543–555.

[21] Monslow MA, Elbashir S, Sullivan NL, et al. Immunogenicity generated by mRNA vaccine encoding VZV gE antigen is comparable to adjuvanted subunit vaccine and better than live attenuated vaccine in nonhuman primates. Vaccine. 2020 Aug 10; 38(36):5793–5802.

[22] Zhao H, Shao X, Yu Y, et al. A therapeutic hepatitis B mRNA vaccine with strong immunogenicity and persistent virological suppression. bioRxiv. 2023:2022.11.18.517095.

[23] Yang R, Deng Y, Huang B, et al. A core-shell structured COVID-19 mRNA vaccine with favorable biodistribution pattern and promising immunity. Signal Transduct Target Ther. 2021 May 31; 6(1):213.

[24] Zhang H, Zhang L, Lin A, et al. Algorithm for optimized mRNA design improves stability and immunogenicity. Nature. 2023 Sep; 621(7978):396–403.

[25] Lin A, Liang F, Thompson EA, et al. Rhesus Macaque Myeloid-Derived Suppressor Cells Demonstrate T Cell Inhibitory Functions and Are Transiently Increased after Vaccination. J Immunol. 2018 Jan 1; 200(1):286–294.

[26] Malavige GN, Jones L, Black AP, et al. Varicella zoster virus glycoprotein E-specific CD4+ T cells show evidence of recent activation and effector differentiation, consistent with frequent exposure to replicative cycle antigens in healthy immune donors. Clin Exp Immunol. 2008 Jun; 152(3):522–31.

[27] Lu LL, Suscovich TJ, Fortune SM, et al. Beyond binding: antibody effector functions in infectious diseases. Nat Rev Immunol. 2018 Jan; 18(1):46–61.

[28] Boeren M, Meysman P, Laukens K, et al. T cell immunity in HSV-1- and VZV-infected neural ganglia. Trends Microbiol. 2023 Jan; 31(1):51–61.

[29] Alameh MG, Tombácz I, Bettini E, et al. Lipid nanoparticles enhance the efficacy of mRNA and protein subunit vaccines by inducing robust T follicular helper cell and humoral responses. Immunity. 2021 Dec 14; 54(12):2877–2892.e7.

[30] Connors J, Joyner D, Mege NJ, et al. Lipid nanoparticles (LNP) induce activation and maturation of antigen presenting cells in young and aged individuals. Commun Biol. 2023 Feb 17; 6(1):188.

[31] Lenart K, Hellgren F, Ols S, et al. A third dose of the unmodified COVID-19 mRNA vaccine CVnCoV enhances quality and quantity of immune responses. Mol Ther Methods Clin Dev. 2022 Dec 8; 27:309–323.

[32] Hellgren F, Cagigi A, Arcoverde Cerveira R, et al. Unmodified rabies mRNA vaccine elicits high cross-neutralizing antibody titers and diverse B cell memory responses. Nat Commun. 2023 Jun 22; 14(1):3713.

[33] Thompson EA, Ols S, Miura K, et al. TLR-adjuvanted nanoparticle vaccines differentially influence the quality and longevity of responses to malaria antigen Pfs25. JCI Insight. 2018 May 17; 3(10).

[34] Jonsson-Schmunk K, Ghose R, Croyle MA. Immunization and Drug Metabolizing Enzymes: Focus on Hepatic Cytochrome P450 3A. Expert Rev Vaccines. 2021 May; 20(5):623–634.

[35] McColl ER, Croyle MA, Zamboni WC, et al. COVID-19 Vaccines and the Virus: Impact on Drug Metabolism and Pharmacokinetics. Drug Metab Dispos. 2023 Jan; 51(1):130–141.

[36] Fiore J, Co-van der Mee MM, Maldonado A, et al. Safety and reactogenicity of the adjuvanted recombinant zoster vaccine: experience from clinical trials and post-marketing surveillance. Ther Adv Vaccines Immunother. 2021; 9:25151355211057479.

[37] Sun HX, Xie Y, Ye YP. Advances in saponin-based adjuvants. Vaccine. 2009 Mar 13; 27(12):1787–96.

[38] Del Giudice G, Rappuoli R, Didierlaurent AM. Correlates of adjuvanticity: A review on adjuvants in licensed vaccines. Semin Immunol. 2018 Oct; 39:14–21.

[39] Chaudhary N, Weissman D, Whitehead KA. mRNA vaccines for infectious diseases: principles, delivery and clinical translation. Nat Rev Drug Discov. 2021 Nov; 20(11):817–838.

[40] Bogger-Goren S, Baba K, Hurley P, et al. Antibody response to varicella-zoster virus after natural or vaccine-induced infection. J Infect Dis. 1982 Aug; 146(2):260–5.

[41] Levin MJ, Weinberg A. Immune responses to zoster vaccines. Hum Vaccin Immunother. 2019; 15(4):772–777.

[42] Tang H, Moriishi E, Okamoto S, et al. A community-based survey of varicella-zoster virus-specific immune responses in the elderly. J Clin Virol. 2012 Sep; 55(1):46–50.

[43] Hata A, Asanuma H, Rinki M, et al. Use of an inactivated varicella vaccine in recipients of hematopoietic-cell transplants. N Engl J Med. 2002 Jul 4; 347(1):26–34.

[44] Jennewein MF, Alter G. The Immunoregulatory Roles of Antibody Glycosylation. Trends Immunol. 2017 May; 38(5):358–372.

[45] Jefferis R. Glycosylation as a strategy to improve antibody-based therapeutics. Nat Rev Drug Discov. 2009 Mar;8(3):226–34.

[46] Kao D, Lux A, Schaffert A, et al. IgG subclass and vaccination stimulus determine changes in antigen specific antibody glycosylation in mice. Eur J Immunol. 2017 Dec; 47(12):2070–2079.

[47] Mahan AE, Jennewein MF, Suscovich T, et al. Antigen-Specific Antibody Glycosylation Is Regulated via Vaccination. PLoS Pathog. 2016 Mar; 12(3):e1005456.

[48] Sei JJ, Cox KS, Dubey SA, et al. Effector and Central Memory Poly-Functional CD4(+) and CD8(+) T Cells are Boosted upon ZOSTAVAX(®) Vaccination. Front Immunol. 2015; 6:553.

[49] Corbett KS, Edwards DK, Leist SR, et al. SARS-CoV-2 mRNA vaccine design enabled by prototype pathogen preparedness. Nature. 2020 Oct; 586(7830):567–571.

[50] Vogel AB, Kanevsky I, Che Y, et al. BNT162b vaccines protect rhesus macaques from SARS-CoV-2. Nature. 2021 Apr; 592(7853):283–289.

[51] Chivukula S, Plitnik T, Tibbitts T, et al. Development of multivalent mRNA vaccine candidates for seasonal or pandemic influenza. NPJ Vaccines. 2021 Dec 16; 6(1):153.

[52] Coccia M, Collignon C, Hervé C, et al. Cellular and molecular synergy in AS01-adjuvanted vaccines results in an early IFNγ response promoting vaccine immunogenicity. NPJ Vaccines. 2017; 2:25.

[53] De Giovanni M, Cutillo V, Giladi A, et al. Spatiotemporal regulation of type I interferon expression determines the antiviral polarization of CD4(+) T cells. Nat Immunol. 2020 Mar; 21(3):321–330.

[54] Liang F, Lindgren G, Lin A, et al. Efficient Targeting and Activation of Antigen-Presenting Cells In Vivo after Modified mRNA Vaccine Administration in Rhesus Macaques. Mol Ther. 2017 Dec 6;25(12):2635–2647.

[55] Nam HJ, Hong SJ, Lee A, et al. An adjuvanted zoster vaccine elicits potent cellular immune responses in mice without QS21. NPJ Vaccines. 2022 Apr 22; 7(1):45.

[56] Haberthur K, Messaoudi I. Animal models of varicella zoster virus infection. Pathogens. 2013 May 13; 2(2):364–82.

