## Supplemental materials for "Herpes zoster mRNA vaccine induces superior vaccine immunity over licensed vaccine in mice and rhesus macaques"

**Supplemental Materials and Methods**

**Antibody Function Assay**

Antibody-dependent complement deposition (ADCD), antibody dependent neutrophil phagocytosis (ADNP), and antibody dependent cellular phagocytosis (ADCP) assays were carried out to assess the functions of antibodies as previously described with minor modifications [1-3]. In brief, VZV gE protein were biotinylated (Frdbio, China) and coupled at a 1:1 ratio to 1-μm red fluorescent neutravidin beads (Thermo Fisher) for 2 h at 37 °C in dark. Excess antigens were removed by washing twice with PBS containing 0.1% BSA. Antigen-coupled beads were then incubated with diluted serum samples for 2 hours at 37°C in dark to facilitate immune complex formation and then washed to remove unbound immunoglobulins. For ADCD assay, guinea pig complements were added to the immune complexes for 15 min at 37 °C followed by washing twice and stained with a fluorescein-conjugated goat anti-guinea pig complement C3 Ab (Invitrogen). For ADNP assay, neutrophils (above 90% purity) isolated from murine bone marrow by percoll density centrifugation were added to the immune complexes and incubated for 2 h at 37 °C. For ADCP assay, 25,000 THP-1 cells were added to the immune complexes and incubated for 8 h at 37 °C. Samples were fixed with 4% paraformaldehyde and acquired on Attune NxT Flow Cytometer (Thermo). All events were gated on single live cells and beads-positive events. For ADCD assay, the mean fluorescence intensity of C3-positive events was reported. For ADNP and ADCP assays, cells were acquired to calculate the phagocytic score. Phagocytic score = % beads positive cells × GMFI of beads positive cells. GMFI denotes geometric mean fluorescence intensity. Data were analyzed using FlowJo V.10.1 (Tree Star).

**Measurement of antigen-specific AIM+ T cells and Tfh cells**

2 million splenocytes were stimulated with or without gE protein (2 μg/sample, Acro Biosystems) for 20 hours. Afterwards, cells were stained with LIVE/DEAD™ Fixable Aqua Dead Cell Stain Kit (Thermo) for 5 minutes and then incubated with antibody cocktails and Fc receptor blocking reagent (Miltenyi) for 20 minutes at 4°C in dark. Flow cytometric analysis was performed on a BD FACSymphony A3 (BD Biosciences). A list of antibodies used in this analysis is available in supplemental materials.

**RNA extraction, library preparation and sequencing**

Total RNAs from rhesus PBMCs isolated 24 hours after boost immunization were extracted using TRIzol Reagent (Invitrogen) based on the methods from reported protocol [4]. After RNA extraction, DNase I was used to digest DNA. The quality of the RNA was determined by A260/A280 with Nanodrop^TM^ One Spectrophotometer (Thermo Fisher Scientific Inc). The integrity of RNAs was confirmed by electrophoresis on 1.5% agarose gels. Quantification of RNAs was performed with Qubit 3.0 with Qubit^TM^ RNA Broad Range Assay kit (Life Technologies). 2 μg of total RNAs was used for the preparation of the stranded RNA sequencing library using the KC-Digital^TM^ Stranded mRNA Library Prep Kit for Illumina® (Wuhan Seqhealth Co., Ltd. China), according to the manufacturer's instructions. The library products (200-500bps) were enriched, quantified, and finally sequenced on DNBSEQ-T7 sequencer (MGI Tech Co., Ltd. China) with PE150 model.

**Transcriptomics analysis**

Raw sequencing data were filtered by Trimmomatic (version 0.36), low-quality reads were discarded, and adaptor sequences-contaminated reads were trimmed. Then, the clean reads were processed with in-house scripts to remove duplication bias introduced during library preparation and sequencing. The de-duplicated consensus sequences were used for standard transcriptomics analysis. The raw sequencing data have been submitted to the NCBI SRA database (accession no. PRJNA1001610). For Venn diagram and volcano plots, differentially expressed genes (DEGs) were identified based on fold-change > 2 or < 0.5 compared to matched pre-vaccination controls at the respective groups, with p-value < 0.05. Clustering and heatmaps are generated using R. Ratios are used to correct gene expression values on Day 29 and Day 0 for each corresponding sample. DEGs were identified with p-value < 0.05 and normalized by z-score normalization. For gene set functional enrichment analysis, we first used the gene GO annotations from the R package org.Hs.eg.db (version 3.1.0) as the background. Genes were mapped to the background set, and the R package clusterProfiler (version 3.14.3) was used for enrichment analysis to obtain the results of gene set enrichment. The minimum gene set was set to 5 and the maximum gene set was set to 5000. Then, for Gene Set Enrichment Analysis (GSEA), samples were compared before and after vaccination. The GSEA software (version 3.0) and the subset (c5.go.bp.v7.4.symbols.gmt) from the Molecular Signatures Database were used to assess relevant pathways and molecular mechanisms. The analysis was based on gene expression profiles and phenotype grouping, with a minimum gene set of 5, a maximum gene set of 5000, and 1000 permutations. The p-value < 0.05 and the FDR < 0.25 were considered statistically significant in GSEA and GO analysis. For clustering of significantly correlated values, we performed correlation analysis on 45 immune-related parameters, which were analyzed using a multivariate nonparametric pearman’s test and 2-tailed P value. The parameters of pathways or blood transcript modules (BTMs) were determined by average fold change value of genes.

**Hematology and blood** **biochemistry**

For hematological evaluation, EDTA-anticoagulated blood was tested on an ADVIA-2120 Hematology Analyzer (Simens, IL). Counts of leukocyte, lymphocyte, monocyte, neutrophil, basophil and eosinophil were measured. For blood biochemical analyses, blood samples were taken into tubes containing the anticoagulant lithium heparin. After centrifugation, the following serum parameters were assayed by Dimension Xpand Plus (Simens): aspartate aminotransferase (AST), blood urea nitrogen (BUN), alanine transaminase (ALT), gamma-glutamyl transpeptidase (GGT), chloridion (CL), total bilirubin (TBI), triglyceride (TGL), creatinine (CREA), plasma albumin (ALB), glucose (GLUC), cholesterol (CHOL), total protein (TP).

**References**

[1]	Fischinger S, Fallon JK, Michell AR, et al. A high-throughput, bead-based, antigen-specific assay to assess the ability of antibodies to induce complement activation. J Immunol Methods. 2019 Oct; 473:112630.

[2]	Karsten CB, Mehta N, Shin SA, et al. A versatile high-throughput assay to characterize antibody-mediated neutrophil phagocytosis. J Immunol Methods. 2019 Aug; 471:46-56.

[3]	Ackerman ME, Moldt B, Wyatt RT, et al. A robust, high-throughput assay to determine the phagocytic activity of clinical antibody samples. J Immunol Methods. 2011 Mar 7;366(1-2):8-19.

[4]	Chomczynski P, Sacchi N. Single-step method of RNA isolation by acid guanidinium thiocyanate-phenol-chloroform extraction. Anal Biochem. 1987 Apr;162(1):156-9.

**Supplemental Figure 1**

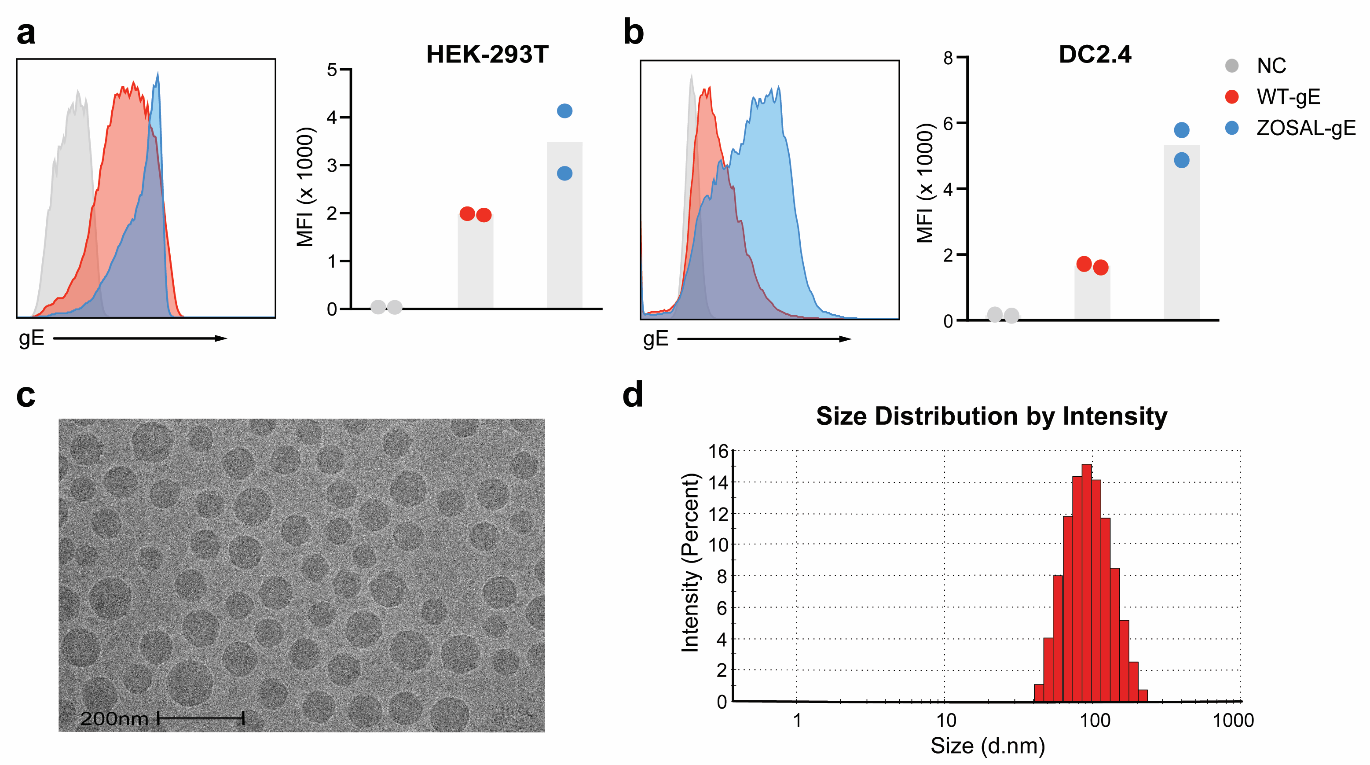

**Figure S1. Characterization of gE-mRNAs and the mRNA vaccine formulation (ZOSAL)**

**a-b.** Protein expression of gE mRNAs was evaluated by flow cytometry upon transfection into HEK293T (**a**) or DC2.4 (**b**) cells for 24 hours. Untreated cells were used as negative control (NC). WT and ZOSAL represent the wide-type and codon-optimized gE-mRNA sequence, respectively. Data from two independent experiments are shown. **c.** Evaluation of ZOSAL by Transmission Electron Microscopy (TEM). Representative image is shown. **d.** Size distribution of ZOSAL with Dynamic Light Scattering (DLS).

**Supplemental Figure 2**

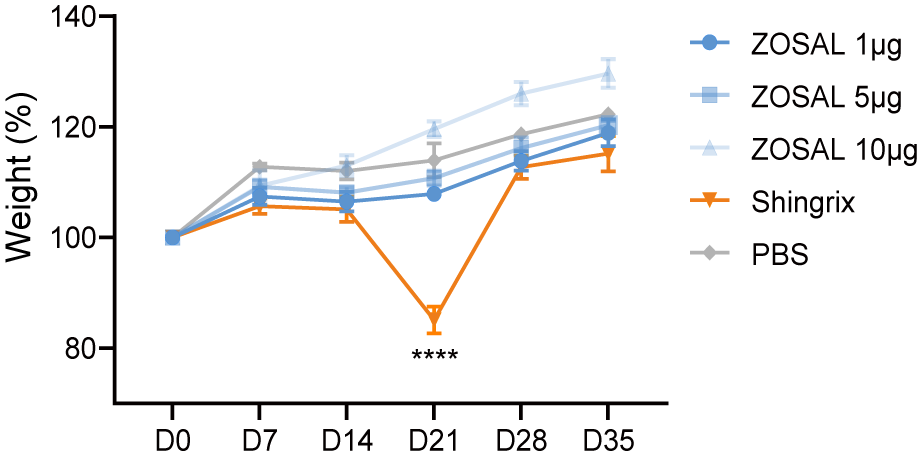

**Figure S2. Monitoring of body weight following vaccination.**

C57BL/6 mice (n=6) were i.m. immunized with escalating doses of ZOSAL or 0.1 human dose of Shingrix on day 0 and day 14. Mice receiving PBS injection were used as control. Body weight was monitored, and percentage of weight change is shown. Statistical difference was assessed using 2-way ANOVA. ****p ≤ 0.001.

**Supplemental Figure 3**

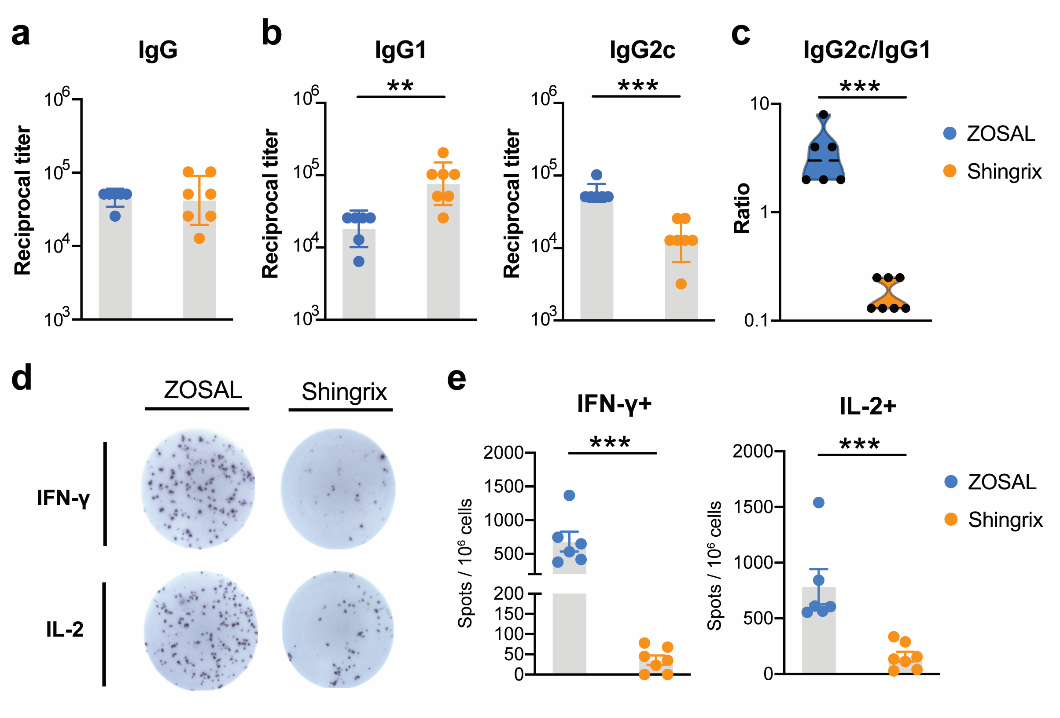

**Figure S3. Evaluation of immunogenicity of ZOSAL and Shingrix in aged mice.**

10-month old C57BL/6 mice (n=6) were immunized i.m. with two doses of 5μg ZOSAL or 0.1 human dose of Shingrix at a 2-week interval. **a-b.** Endpoint titers of anti-gE IgG, IgG1 and IgG2c at day 14 after the boost immunization were measured by ELISA. **c.** Ratio of IgG2c/IgG1 is shown. **d-e.** Frequencies of IFN-γ or IL-2-secreting T cells were measured by ELISpot. Data are shown as mean ± SEM. Mann-Whitney U test was used for statistical analysis. **p ≤ 0.01, ***p ≤ 0.001.

**Supplemental Figure 4**

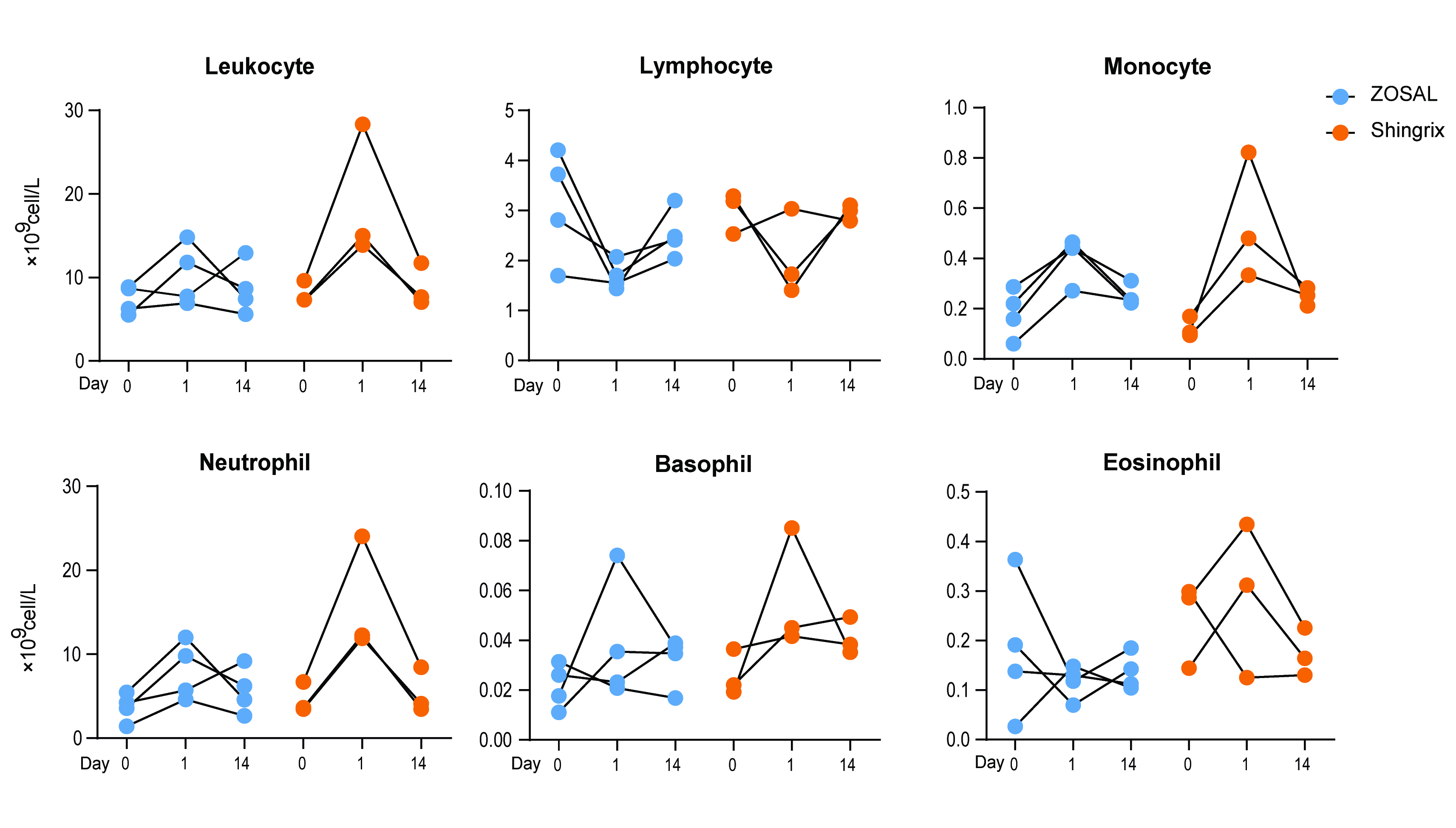

**Figure S4. Measurement of complete blood counts in vaccinated rhesus macaques.** Rhesus macaques were immunized i.m. with ZOSAL (n=4) or Shingrix (n=3) at an interval of 4 weeks. Complete blood counts were measured from whole blood collected before vaccination and at day 1, 14 after prime vaccination.

**Supplemental Figure 5**

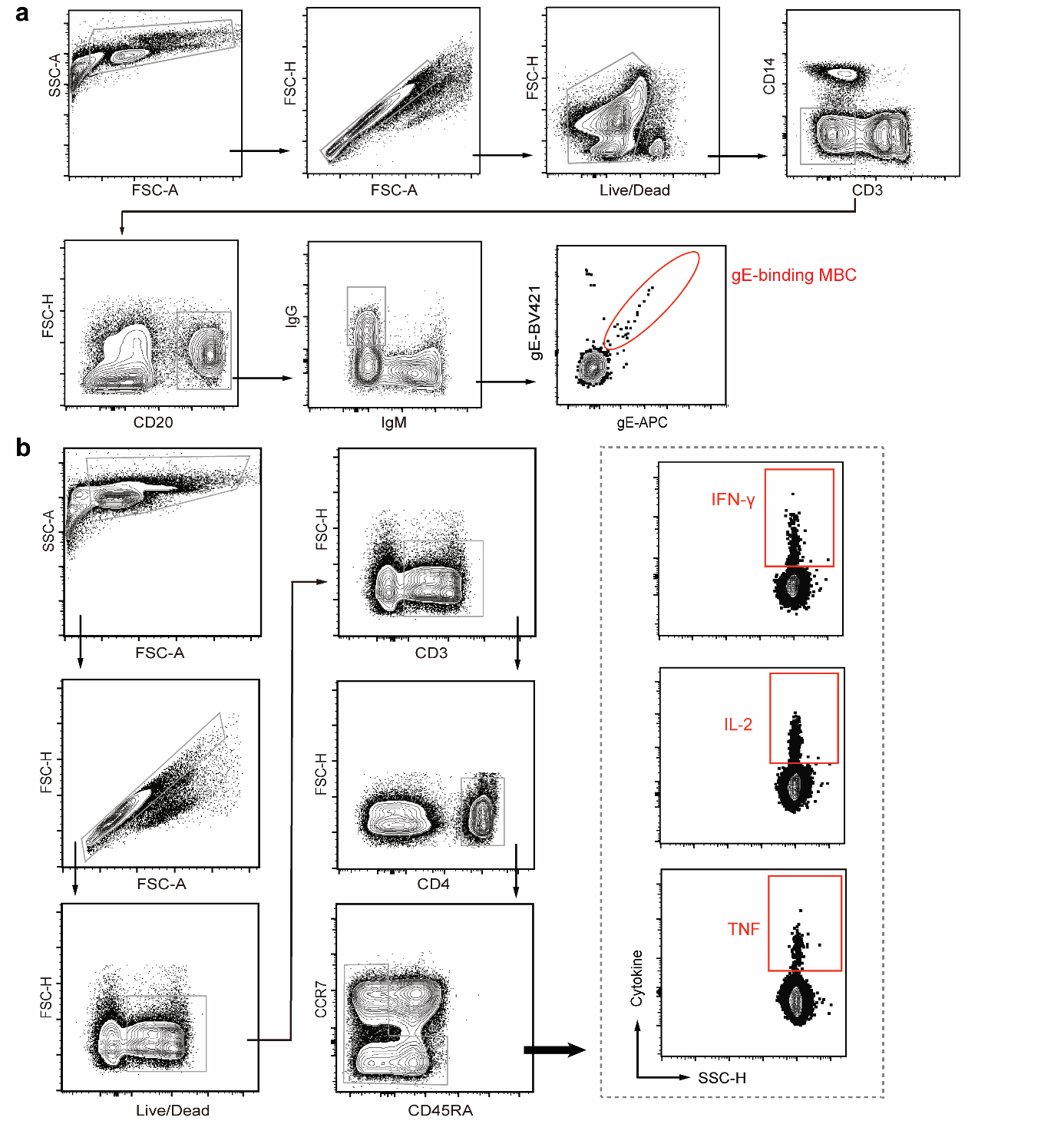

**Figure S5. Representative gating strategy for flow cytometry analysis of MBC and T cell response in rhesus macaques.**

**a**. Gating strategy for phenotypic identification of class-switched gE-binding MBC in blood of rhesus macaques. Data from one representative animal is shown. **b**. Gating strategy for analysis of IFN-γ, IL-2, or TNF-secreting CD4^+^ T cells in blood of rhesus macaques. Data from staphylococcus enterotoxin B stimulated PBMCs is used as an example to show.

**Supplemental Tables**

Table S1. List of anti-mouse antibodies used for FACS analysis

| **Antibody** | **Clone** | **Manufacturer** |
| --- | --- | --- |
| IgM | RMM-1 | Biolegend |
| IgD | 11-26c.2a | Biolegend |
| CD45R/B220 | RA3-6B2 | Biolegend |
| CD19 | 6D5 | Biolegend |
| CD38 | 90 | Biolegend |
| CD3 | 17A2 | Biolegend |
| CD4 | GK1.5 | Biolegend |
| CD44 | IM7 | Biolegend |
| CD62L | MEL-14 | Biolegend |
| IFN-γ | XMG1.2 | Biolegend |
| TNF | MP6-XT22 | Biolegend |
| IL-2 | JES6-5H4 | Biolegend |
| IL-21 | mhalx21 | Invitrogen |
| CD4 | RM4-5 | Biolegend |
| CXCR5 | L138D7 | Biolegend |
| PD1 | 29F.1A12 | Biolegend |
| ICOS | C398.4A | Biolegend |
| OX40 | OX-86 | Biolegend |
| CD137 | 17B5 | Biolegend |
| Gr-1 | RB6-8C5 | Biolegend |

Table S2. List of anti-rhesus macaque antibodies used for FACS analysis

| **Antibody** | **Clone** | **Manufacturer** |
| --- | --- | --- |
| CD3 | SP34-2 | BD |
| CD8 | RPA-T8 | BD |
| CD20 | 2H7 | BD |
| CD14 | M5E2 | Biolegend |
| CD16 | 3G8 | Biolegend |
| HLA-DR | L243 | Biolegend |
| IgM | MHM-88 | Biolegend |
| IgG | G18-145 | BD |
| CD20 | 2H7 | BD |
| CD4 | L200 | BD |
| CD45RA | 5H9 | BD |
| CCR7 | G043H7 | Biolegend |
| IL-2 | MQ1-17H12 | Biolegend |
| IFN-γ | 4S.B3 | Biolegend |
| TNF | MAb11 | Biolegend |
